# A regulator controls both DNA damage response and anti-phage defense networks in *Moraxellaceae*

**DOI:** 10.1101/2024.11.11.622897

**Authors:** Shuang Song, Shitong Zhong, Qiucheng Shi, Xiangkuan Zheng, Yue Yao, Wenxiu Wang, Shanhou Chen, Zijun Huang, Dongyue An, Hong Xu, Bing Tian, Ye Zhao, Liangyan Wang, Wei Zhang, Xiaoting Hua, Yunsong Yu, Huizhi Lu, Lu Fan, Yuejin Hua

**Author notes:** These authors contribute equally to this work (Shuang Song, Shitong Zhong). To whom correspondence should be addressed.; Email: H. Y., Correspondence may also be addressed to, or.

## Abstract

DNA damage chemicals, including many antibiotics, often induce prophage induction and phage outbreaks in microbial communities, a major threat to bacterial survival. *Moraxellaceae* contains clinically highly relevant strains with outstanding antibiotic and radio resistance, yet their cellular-level regulation in DNA damage response and anti-phage defense is largely unknown. Here, we identified a WYL family protein that had replaced the ubiquitous SOS system in evolution and directly regulated a transcriptional network of both DNA repair genes and anti-phage defense genes under DNA damage stress. This study depicts a mechanism that how bacteria maintain immunity to phages without compromising on antibiotic resistance, and sheds light on controlling the resistance of *Moraxellaceae* strains in clinical practice.

## Main

The family *Moraxellaceae* is composed of a highly diverse group of bacteria with clinical significance^1^, including some lineages as human pathogens with multidrug-resistance like *Acinetobacter baumannii*^2^ and others with radio-resistance like *Acinetobacter radioresistens*^3,4^. Up to 70% mortality rate caused by antibiotic-resistant *A. baumanii* strains has been reported^5^. Many antibiotics target DNA or DNA replication pathways in bacterial cells and have overlapping action modes with common DNA damage agents such as mitomycin C (MMC), cisplatin, and UV radiation^6^. It is a long-standing question of how bacteria of *Moraxellaceae* respond to DNA damage^7,8^, as they encode none of the three known bacterial global DNA damage response (DDR) systems, namely SOS^9^, PprI-DdrO^10^, and PafBC^11^. While the protein UmuDAb was found to directly regulate three DNA damage-induced genes including itself in the genus *Acinetobacter*^12–14^, knockout of *umuDAb* was insufficient to cause a decline in the survival rate of *A. baumanii* under UV radiation^15,16^. Therefore, whether there is any global DDR system in *Moraxellaceae* is still unknown^17^.

Prophages are the dormant state of lysogenic phages and are often integrated into the host chromosomes^18^. Stimulated by environmental stresses including DNA damage agents such as UV^19^, MMC^17^, and colibactin^20^, the activation of prophage proteins facilitates prophage genome excision and replication and eventually leads to the lysis of host cells^21^. It has been frequently reported in bacteria including *Moraxellaceae* strains that prophages were induced by DNA damage agents^9,17^, introducing an additional threat to the cells^20,22^. Moreover, the induced phages may infect closely related strains in the same habitat, resulting in a phage epidemic^23^. Bacteria have developed multiple antiviral strategies such as the restriction-modification (RM) systems, the clustered regularly interspaced short palindromic repeats–CRISPR-associated proteins (CRISPR–Cas) systems, and Zorya systems^18,24^. Whereas the upregulation of Cas genes in DNA damage conditions has been reported in *Acinetobacter* strains^17,25^, implying a possible connection between DNA damage and anti-phage defense responses, how bacteria cope with prophage induction after DNA damage remains largely unknown.

In this study, we aim to identify the global regulatory network for DDR in *Moraxellaceae*. The binding motif of this potential system was identified by applying a comparative genomics strategy. DNA pulldown assays excavated out a newly classified WYL family protein binding to this motif, whereas evolution analyses and transcriptomic experiments verified that this protein functions as a transcriptional activator for both conventional DDR and anti-phage defense in the family *Moraxellaceae*. The knockout of the protein caused a severe decrease in the survival rate of *A. baumannii* under DNA damage and phage infection treatments, verifying a unique DDR mechanism regulated by this protein in those highly resistant strains of *Moraxellaceae*.

## Results

### A conserved motif is involved in a putative global DDR system in *Acinetobacter*

We compared the promoter regions of DDR genes to identify potential global DDR regulation systems in *Acinetobacter*, a major genus of *Moraxellaceae* with well noted resistance. Based on previously published transcriptomic data^17,26^ (Supplementary Table 1), the promoter regions of 63 genes likely involved in DDR were analyzed in 88 representative *Acinetobacter* genomes (Fig. 1a, Supplementary Table 2). A conserved and well-palindromic DNA motif was identified appearing in the promoter region of more than 20 genes including *recA* for homologous recombination DNA damage repair, *aciT* for ciprofloxacin tolerance, *ssb* for single-stranded DNA protection, *uvrA* and *uvrC* for nucleotide excision repair, *gyrA* and *gyrB* for forming DNA supercoils, and dGTPase-*ruvA-ruvB* operon for holiday junction processing, indicating the existence of an undiscovered global DDR system in *Acinetobacter* (Fig. 1b, Supplementary Table 3). In addition, some Cas genes also had this motif in their promoter regions, suggesting anti-phage defense may be involved in this system (Supplementary Table 3).

**Fig. 1.**
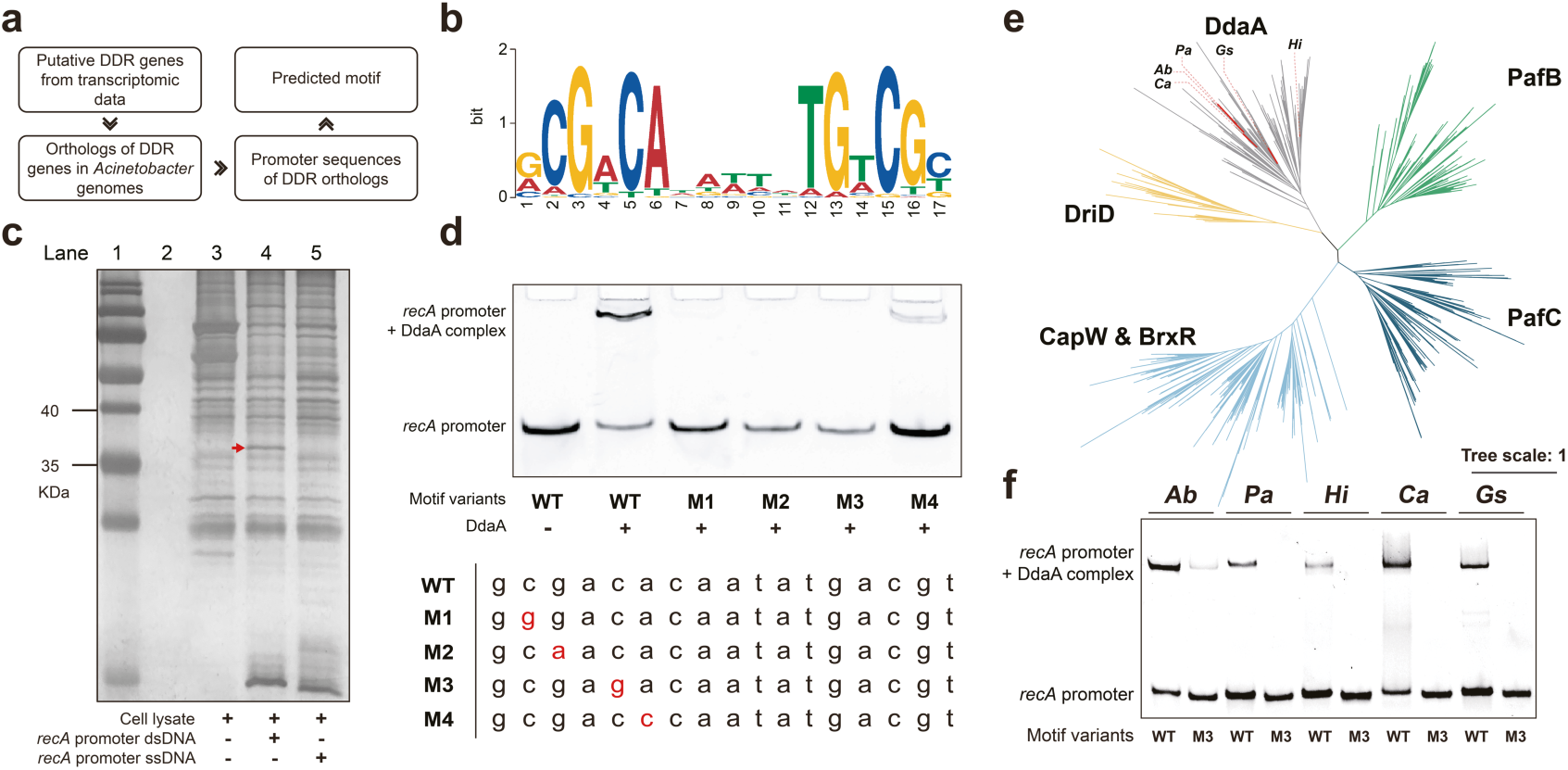
| Discovery of the regulator of a putative global DDR system and its binding motif in *Acinetobacter*. **a**, The schematic bioinformatic workflow to discover the common binding motif in the promoter regions of DDR genes in *Acinetobacter*. **b**, The binding motif of the potential regulator of a global DDR system in *Acinetobacter*. **c**, DNA pull-down assay of *A. radioresistens* ATCC 43998 with *recA*’s promoter. The protein marker is PageRuler™ 26616 in lane 1. Lane 2 is a blank. Lane 3 is the negative control, where no DNA was added. Lane 4 is the dsDNA pulldown result. Lane 5 is the ssDNA pulldown result. The red arrow shows a protein band that only appears in Line 4. **d**, EMSA of DdaA with *recA*’s promoter motif of *A. radioresistens* ATCC 43998. The mutated bases are highlighted in red. **e**, The maximum-likelihood tree of WYL domain-containing proteins. Each cluster is shown in different colors. Branches of DdaA homologs are in gray and those validated by EMSA as shown in panel **f** are in red. Ab = *A. baumannii* ATCC 17978. Pa = *Psychrobacter aquimaris* KCTC 12254. Hi = *Haemophilus influenzae* 65290_NP_Hi3. Ca = *Chrysiogenes arsenatis* DSM 11915. Gs = *Geoalkalibacter subterraneus* Red1. **f**, EMSA of DdaA homologs and the promoter sequence of *recA* from *A. radioresistens* ATCC 43998.

### A potential regulatory protein specifically binds to this motif

To identify the regulator of this putative DDR system, we conducted a pulldown experiment by incubating the cell lysate of the radio-resistant strain *A. radioresistens* ATCC 43998 with beads coated with the promoter DNA of its *recA* gene. Followed by electrophoretic separation of the enriched proteins, a protein band appeared in the 35 to 40 kDa size range in the lane, where the double-strand DNA (dsDNA) of the promoter was used, but not in the lane, where the single-strand DNA (ssDNA) of the promoter was used or in the negative control lane, where no DNA was added (Fig. 1c). The mass spectrometry result indicates that this protein contains a WYL and a HTH domain with a molecular weight of 38.685 kDa matching UniProt Accession ID: A0A7U9PMQ2 (Supplementary Fig. 1ab, Supplementary Table 4). As this protein is possibly involved in regulating DDR genes in *Acinetobacter*, we name it ‘DNA damage response protein *Acinetobacter* A’ (DdaA) hereafter.

The specific interaction between the protein and the motif was further confirmed by in vitro electrophoretic mobility shift assay (EMSA) with DdaA from *A. radioresistens* and motif-containing promoter region of *recA*. The band of DdaA binding to an intact motif shows a significant mobility shift (Fig. 1d). Mutations of any of the four conserved base pairs in the motif drastically impaired the binding, verifying that these sites may be specifically involved in DdaA binding. The M4 mutant retained a weak shifted band, indicating that this site is less essential than the other three.

### DdaA homologs form a major cluster in the WYL protein family

WYL domain-containing proteins are ubiquitous in bacteria and function as regulators in bacterial immunity^27–29^ or DDR^11,30^. Phylogenetic analysis indicates that homologs of DDR regulators DdaA, DriD, PafB, and PafC, respectively, form four independent monophyletic clusters, while homologs of anti-phage defense regulators CapW and BrxR form another monophyletic cluster (Fig. 1e). Their neighboring positions in the tree and the fact that their binding motifs are similar suggest that DdaA and DriD may share a common ancestor (Fig. 1e, Supplementary Text, Supplementary Fig. 1cd). Like other class A WYL domain-containing proteins, DdaA consists of an N-terminal wHTH domain, a WYL domain, and a WCX (WYL protein C-terminal extension) domain^31^, and it forms a homodimer in the solution (Supplementary Text, Supplementary Fig. 2).

To investigate the conservation of DdaA in binding to the identified motif, homologs sharing 84%, 45%, 38%, 34%, and 29% identities with the DdaA of *A. radioresistens* ATCC 43998 from five strains, including *A. baumannii* ATCC 17978, *Psychrobacter aquimaris* KCTC 12254, *Haemophilus influenzae* strain 65290_NP_Hi3*, Chrysiogenes arsenatis* DSM 11915, and *Geoalkalibacter subterraneus* Red1 (Fig. 1e), were purified and subjected to EMSA with the promoter DNA of *recA* from *A. radioresistens* ATCC 43998 containing the motif. All these proteins bond to the motif and mutations on the M3 sites cause a weakened or lost binding capacity of these DdaA homologs (Fig. 1f).

### DdaA regulation replaced the SOS system for global DDR in the early divergence of *Moraxellaceae*

Comparison between the species tree of *Moraxellaceae* and the gene tree of DdaA suggests the gradual establishment of the DdaA-based DDR regulation system in the early divergence of *Moraxellaceae*. The species tree of *Moraxellaceae* and neighboring taxonomic lineages were extracted from the GTDB bacterial tree for analysis (Supplementary Fig. 3). A copy of *ddaA* gene (DdaA-1) was horizontally transferred from a gammaproteobacterium to the common ancestor of *Alcanivoracaceae* and *Moraxellaceae* (Fig. 2a, Supplementary Fig. 4, Supplementary Fig. 5, Supplementary Text). Before this transfer, DdaA was a putative regulator protein of various genes located near the *ddaA* gene locus (Supplementary Fig. 4b). However, the function of these regulated genes is currently unknown. After being obtained, this copy of DdaA was vertically inherited in the two families of *Alcanivoracaceae* and *Moraxellaceae* (Fig. 2a, Supplementary Fig. 5). However, in *Alcanivoracaceae*, in the basal lineages of *Moraxellaceae* and the genus *Moraxella*, the *ddaA* gene was sporadically lost and often replaced by copies of DdaA horizontally transferred from various bacteria (DdaA-other) (Fig. 2a, Supplementary Fig. 4a and Supplementary Fig. 5).

**Fig. 2.**
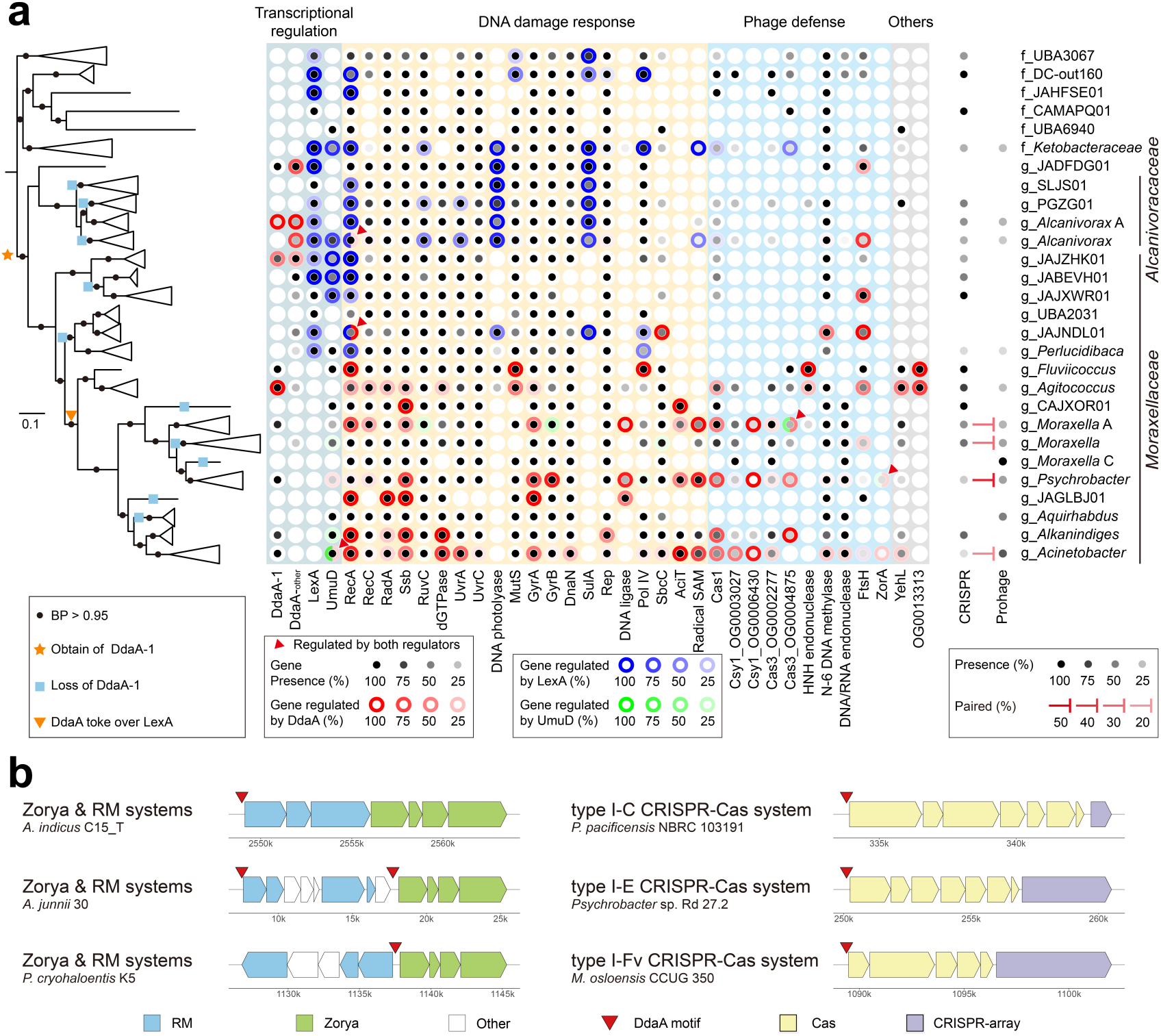
| The distribution and evolutionary route of DdaA, LexA, UmuD and the genes they regulate in *Moraxellaceae*. **a**, The phylogenomic maximum-likelihood tree of species in *Moraxellaceae* and neighboring taxonomic lineages is shown on the left panel. Branches are collapsed at the genus level, except in the outgroups at the family level based on GTDB taxonomy^57^. The IQ-Tree ultrafast bootstrapping values > 95% are indicated by black dots on branches. Evolutionary events of DdaA-1 are labeled on branches. In the middle panel, the darkness of the black dots shows the percentage of genomes in the taxonomic clade encoding the protein. The darkness of the colored ring around a dot indicates the percentage of this gene is regulated by DdaA (red), LexA (blue) or UmuD (green) in the taxonomic clade. The matching between pairs of CRISPR-Cas spacers and protospacers on prophages were shown on the right panel. The darkness of the black dots shows the percentage of genomes encoding CRISPR-Cas or prophages. The darkness of the T bars shows the percentage of CRISPR-Prophage pairs in total genomes in the genera. **b**, Representative cases of the locations of DdaA binding motifs in the promoter region of Zorya, RM, and subtypes of CRISPR-Cas systems in *Moraxellaceae*.

Both competition and complementary between DdaA regulation and SOS system in controlling DDR genes are observed in *Alcanivoracaceae* and the basal lineages of *Moraxellaceae*. In these taxa, LexA regulates essential DDR genes forming typical SOS systems as found in most bacteria (Fig. 2a, Supplementary Fig. 5). In contrast, DdaA often regulates a small number of DDR- and anti-phage defense-related genes including *recA*, *recC*, *sbcC*, N-6 DNA methylase gene, and *ftsH*. Interestingly, the essential gene *recA* was regulated simultaneously by LexA and DdaA in some organisms showing an evolutionarily transition state, when the promoter regions of *recA* can be bound by both regulators^32,33^ (Fig. 2a). This observation suggests that the competition of these two regulators in controlling a global DDR system might have started from the regulation of *recA* gene.

It was not until the common ancestor of *Fluviicoccus*, *Acinetobacter*, *Psychrobacter*, and all other genera in that monophyletic clade, that the role of LexA as a global DDR regulator was completely taken over by DdaA (Fig. 2a, Supplementary Fig. 5). DdaA can regulate up to 20 genes involved in DDR and anti-phage defense such as the RM system, Zorya system, and many subtypes of CRISPR-Cas systems (Fig. 2b, Supplementary Fig. 5). The collection of genes regulated by DdaA varies between genus-level subgroups. For example, *gyrB*, a DNA gyrase subunit, is regulated by DdaA in *Psychrobacter* but not in most *Acinetobacter*, while *recA* and the *dGTPase*-*ruvA*-*ruvB* operon are regulated in the latter but not in the former. Notably, in the basal clade including *Agitococcus lubricus* and *Fluviicoccus keumensis*, DdaA is self-regulated. The self-regulation stopped after the common ancestors of three well-developed genera of *Acinetobacter*, *Moraxella,* and *Psychrobacter*, when the DdaA regulatory network evolved as it recruited many more DDR genes into its regulon, and thereafter became a global regulator of the DDR system.

A decade ago, Hare et al. reported that the UmuD protein is involved in the error-prone DNA replication process in *Acinetobacter*^17^ by regulating itself and UmuC^4^. We found that UmuD homologs were frequently found in lineages of *Moraxellaceae* (Fig. 2a, Supplementary Fig. 5,6, Supplementary Text). Most of them are self-regulated, but some others are regulated by DdaA. Therefore, the UmuD regulator may act as a small complement of the DdaA network in *Acinetobacter,* or function independently as found in *A. baumannii* ATCC 17978.

### DdaA activates the transcription of DDR genes triggered by DNA damage agents

To validate that DdaA is crucially involved in DDR without the potentially complex interference of phage infection, we conducted phenotype assays using *A. baumannii* ATCC 17978, which does not contain known RM or CRISPR-Cas systems (Supplementary Table 5). Other potential anti-phage defense genes in this strain are not regulated by DdaA. The successful reconstructions of the *ddaA*-knockout and compensated strains were validated by DNA sequencing (Supplementary Fig. 7a). After a snap exposure to 5-20 J/m^2^ UV radiation, the viability of the *ddaA*-knockout strain decreased by 10 to 10,000 folds compared with the wild-type or the complemented strain (Fig. 3a, Supplementary Fig. 7b). Moreover, the viability decreased by about 100-folds under 1 μg/mL, 2 μg/mL, or 4 μg/mL MMC treatments for 30 min (Fig. 3b, Supplementary Fig. 7c). Both assays demonstrate the indispensable role of DdaA in DNA damage response.

**Fig. 3.**
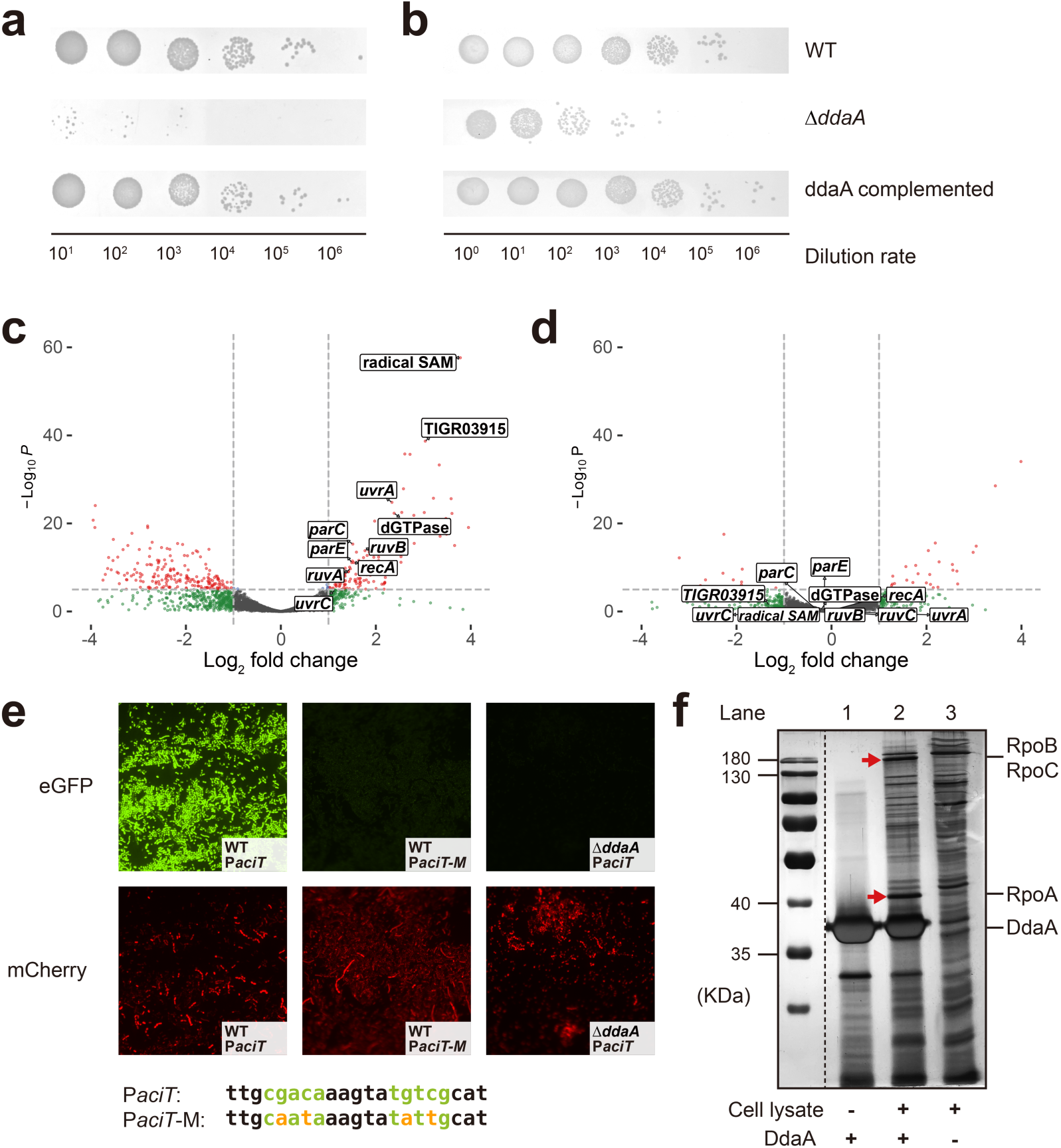
| The activation effect of DdaA on DDR. **a**, Viability of *A. baumannii* ATCC 17978-derived strains after treatment with 10 J/m^2^ UV radiation. **b**, Viability of *A. baumannii* ATCC 17978-derived strains after treatment with 1 μg/mL MMC. **c** and **d,** Volcano plots showing the transcriptional level changes of genes in the wild-type strain (**c**) and *ddaA*-knockout strain (**d**) of *A. baumannii* ATCC 17978 after 15 J/m^2^ UV radiation. The average values of three biological replications are shown. Genes potentially regulated by DdaA are labeled. **e**, Validation of DdaA as a transcriptional activator in *A. baumannii* ATCC 17978-derived strains. P*aciT* = the wild-type strain containing the intact promoter of *aciT*. P*aciT*-M = the wild-type strain harboring a mutated promoter of *aciT*. Green and orange letters show the DdaA binding motif and the mutated bases, respectively. **f**, DNA-DdaA complex pull-down assay. The protein marker is PageRuler™ 26616. Lane 1 is the negative control to determine the non-specific binding by residual proteins in the purification product. The promoter DNA of *recA* of *A. radioresistens* DSM 6976 and the DdaA purification product of *A. baumannii* ATCC 17978 were mixed and added. Lane 3 is the negative control to determine the non-specific DNA binding proteins from *A. baumannii* cell lysate, in which the promoter DNA of *recA* of *A. radioresistens* DSM 6976, and the cell lysate of *A. baumannii* ATCC 17978 were mixed and added. The red arrows suggest the protein bands that only show up in lane 2. In lane 2, the promoter DNA of *recA* from *A. radioresistens* DSM 6976, the DdaA purification product and cell lysate of *A. baumannii* ATCC 17978 were mixed and added to determine the protein(s) potentially interacting with DdaA.

Transcriptional analysis suggests that DdaA is a transcriptional activator of DDR genes. In the *ddaA*-knockout strain of *A. baumannii* ATCC 17978, the basal transcriptions of DDR genes, such as *aciT*, dGTPase, *ruvB*, *uvrA*, the gene of radical SAM protein, and the gene of TIGR03915-family protein, decreased significantly (> 2 folds, *P* < 0.005) compared with the wild-type strain (Supplementary Table 6 and 7), indicating that DdaA is a transcription activator. After exposure to 15 J/m^2^ UV radiation, 118 genes of the wild-type strain were upregulated and 210 were down-regulated (Fig. 3c, Supplementary Table 6). 10 of the upregulated genes contained the conserved motif and were potentially regulated by DdaA. In contrast, only 29 genes were upregulated, and 15 were repressed in the *ddaA*-knockout strain after UV treatment (Fig. 3d, Supplementary Table 6). The transcriptional levels of the *ddaA* gene itself remained unchanged after the UV radiation in this study (Supplementary Table 6), and after MMC^17^ and ciprofloxacin^25^ treatments in previous studies, indicating the activation of DDR genes is not caused by the increased expression level of DdaA. The reverse-transcription quantitative PCR (RT-qPCR) experiment supported the transcriptome results (Supplementary Text, Supplementary Fig. 7d).

To further verify that DdaA is a transcriptional activator *in vivo*, we designed a vector containing the promoter of *aciT* from *A. baumannii*, which harbors the conserved motif bound by DdaA, followed by the gene of eGFP (Fig. 3e). A vector containing a mutated DdaA binding site was constructed for comparison. As a positive background control, these two vectors also contained the promoter of *ompA*, which is not regulated by DdaA but expressed consistently, followed by the gene of mCherry. The first vector was transferred to *A. baumannii* ATCC 17978 wild-type and *ddaA*-knockout strains, while the second was transferred to the wild-type strain. As expected, the expression of mCherry was stable and showed no significant difference between transformed strains (Fig. 3e). The strong expression of the eGFP was detected only in the wild-type strain containing the intact promoter of *aciT*. In contrast, no eGFP signal was detected in the wild-type strain harboring a mutated promoter of *aciT* or the *ddaA*-knockout strain with the intact promoter of *aciT*. This result proves that both DdaA and its binding site are needed to activate the transcription of downstream genes in *A. baumannii*.

WYL domain-containing proteins such as PafBC^11^ and DriD^34^ recruit RNA polymerases (RNAPs) to activate transcription. To explore whether DdaA is involved in RNAP recruitment, we used the complex of DdaA binding to the *recA* promoter of *A. radioresistens* ATCC 43998 as the bait to pull down the protein interacting with this complex from the cell lysate of *A. baumannii*. Two extra bands with molecular weights of around 40 kDa and 180 kDa, respectively, showed up by comparing with the negative controls (Fig. 3f, Supplementary Table 8). Mass spectrometry analysis indicated that the former was RNAP subunit alpha RpoA (37.2kDa, UniProt: A0A828SFM7), and the latter was RNAP subunit beta RpoB (151.8kDa, UniProt: A0A828SUQ7) and RNAP subunit beta RpoC (154.1kDa, UniProt: A0A828SPY9). This result indicates that, like other WYL family proteins, DdaA may also recruit RNAP to activate the transcription of downstream genes.

### DdaA enhances the immunity of *A. baumannii* strains in DNA damage stress

In the DefenseFinder RefSeq database, we found that 90.3% of the CRISPR-Cas system and 27.2% of the RM systems encoded in the *Acinetobacter* are regulated by DdaA. To validate that the control of anti-phage defense genes by DdaA increases the survival rate of *Moraxellaceae* during phage infection, we transferred a vector containing the type I RM system from *A. baumannii* ATCC 19606 with its native promoter harboring the DdaA binding motif (Fig. 4a) to the wild-type *A. baumannii* ATCC 17978, which lacks RM or CRISPR-Cas systems (Supplementary Table 5). Whereas the wild-type *A. baumannii* ATCC 17978 was sensitive to infection of phage PhAb24, the addition of the RM system enhanced its resistance to phage infection (Fig 4b,c). The anti-phage defense ability of the transformed strain was even stronger when treated with DNA damaging agent MMC before phage infection, which promoted the activation effect of DdaA resulting in increased expression of RM genes (Fig. 4d). In contrast, when the *ddaA* gene of *A. baumannii* ATCC 17978 was knocked out or the DdaA binding motif in the promoter region of the RM system was mutated, the gene transcription of the RM systems decreased and the protective effect against phage was negligible either with or without the pretreatment of MMC (Fig. 4b-d). This result confirms that both a functional DdaA and its intact binding motifs in the promoters of anti-phage defense genes are needed to provide defense against phage infection, and this protection mechanism is much more efficient in DNA damage conditions. Moreover, the previous transcriptome analysis of *A. baumannii* when treated with ciprofloxacin shows that the DdaA network is activated under ciprofloxacin pressure^25^. We found that pRM strain’s phage resistance to PhAb24 significantly increased under the 0.2 μg/mL ciprofloxacin treatment (Supplementary Fig. 8a).

**Fig. 4.**
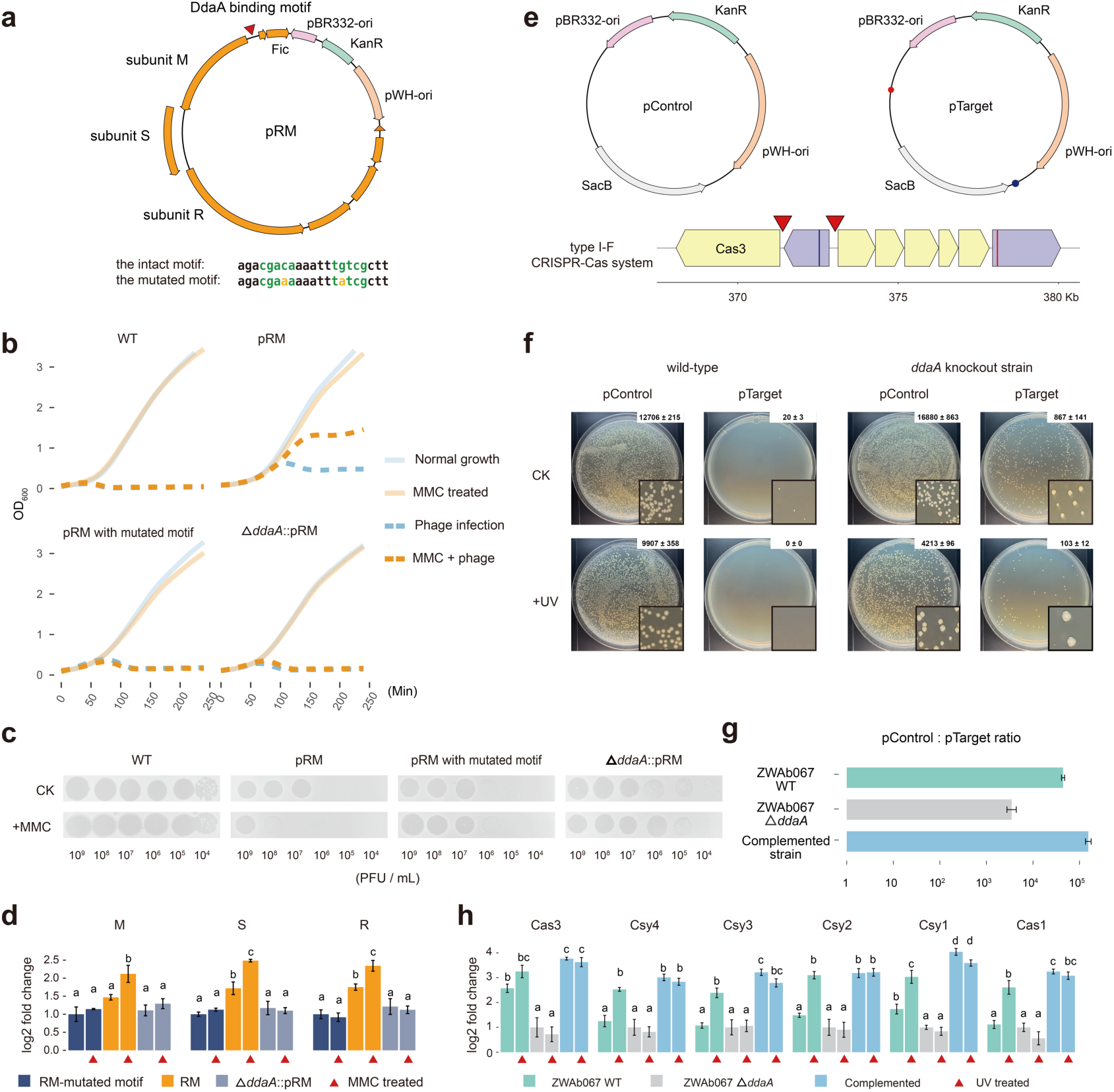
| Validation for the regulatory role of DdaA in anti-phage defense. a, A schematic diagram showing the design of the plasmid pRM. The RM system and its neighboring genes forming a defense locus from *A. baumannii* ATCC 19606 are shown as orange arrows. **b** and **c**, The growth curve (**b**) and phage plaque forming assays (**c**) obtained from co-cultures of *A. baumannii* ATCC 17978-derived strains and phage PhAb24. **d,** The RT-qPCR result of *A. baumannii* ATCC 17978-derived strains. To normalize the data of each gene, the transcription of RM-mutated motif strain was set to 1. **e,** A schematic diagram of the plasmids pTarget and pControl. pControl contains no sequence recorded by CRISPR-array, whereas pTarget contains the protospacer-adjacent motif (2 bp) and protospacer (32 bp) sequences of the prophages from *A. baumannii* strain AB046 (blue dot) and *Acinetobacter pittii* strain FDAARGOS 1397 (red dot). **f,** The plasmid interference assay testing the exogenous DNA removal ability by the native CRISPR-Cas system in *A. baumannii* ZWAb067. Parts of the plate images have been enlarged 25 times to show the size difference between the colonies. **g**, The relative abundance of the pControl versus the pTarget plasmids in overnight cultures of *A. baumannii* ZWAb067 assessed by qPCR. **h**, The RT-qPCR result of *A. baumannii* ZWAb067-derived strains. To normalize for each gene, the transcription levels of *ddaA*-knockout strains are set to 1. Grouping results of the least significant difference (LSD test, alpha=0.05) are labeled for panels **d** and **h**. If two groups share the same small letter, the differences between these groups are not statistically significant.

Many species of *Moraxellaceae* adopting the DdaA global system also have CRISPR-Cas systems and/or prophages (Fig. 3). The CRISPR spacer sequences are frequently found targeting the prophage sequences in the same strain or other strains in the same genus (Fig. 3, Supplementary Fig. 5, Supplementary Table 9). The cross-immunity between *Moraxellaceae* strains indicates their putative cohabitation in the environments. In comparison, cross-immunity is not detected in the basal lineages of *Moraxellaceae* nor in the family of *Alcanivoracaceae*, where the SOS system is dominant.

To mimic the invasion of DNA-damage-induced prophages, we conducted a plasmid interference assay. We designed and transferred a pTarget plasmid to an *A. baumannii* strain ZWAb067 that naturally harbors a functional CRISPR I-F system flanked by two DdaA binding motifs (Fig. 4e). pTarget contains two protospacer sequences specifically targeted by two of the spacers of the CRISPR-Cas I-F system. For comparison, a plasmid without this protospacer sequence namely pControl was used as a control. Successful transformation by these plasmids and evasion of degradation by the native CRISPR-Cas system would enable the bacteria to form colonies on kanamycin-resistant plates. Our results show that in the wild-type strain, the transformation efficiency of pTarget was 0.16% of pControl indicating a functional CRISPR-Cas system (Fig. 4f). In the *ddaA*-knockout strain, this ratio increased to 5.14% suggesting that the deactivation of DdaA impaired the immune efficiency of the CRISPR-Cas system. The complemented strain proved the activation role of DdaA in the CRISPR-Cas system. All pTarget plasmids were immunized and no successful transformation was detected. After exposure to 5 J/m^2^ UV radiation, wild-type cells also achieved a 100% removal rate of pTarget, whereas the *ddaA*-knockout strain remained at 97.5%, suggesting the UV radiation-induced DNA damages further stimulated the activation effect of DdaA to the CRISPR-Cas system (Fig. 4f). We also transferred the mix of pControl and pTarget plasmids at a 1:1 ratio to the ZWAb067-derived strains and tested the relative abundance of pControl to pTarget in overnight cultures by using quantitative PCR (qPCR). The higher the pControl to pTarget ratio, the better the efficiency of the CRISPR-Cas system to remove pTarget plasmids. The complemented strain showed the highest pControl to pTarget ratio at 1.55×10^5^, whereas the *ddaA*-knockout strain showed the lowest at 3.47×10^3^ (Fig. 4g). These results further support that the knockout of the *ddaA* impaired the exogenous DNA removal ability of the CRISPR-Cas system. Furthermore, the RT-qPCR analysis revealed that the knockout of the *ddaA* led to significant reductions in the transcription of Cas genes under normal conditions or after UV radiation (Fig. 4h). These findings collectively suggest that DdaA plays a crucial role in regulating the expressions of Cas genes, particularly in response to DNA damages.

On the other hand, while the knockout of DdaA reduced the basal immune effect of the anti-phage defense systems in *A. baumannii* strains (Fig. 4), revisiting previously published transcriptomic data^35^ shows no up-regulation of DdaA-controlled genes during phage infection (Supplementary Fig. 8b). The transcription of DdaA regulated gene in *A. baumannii* ATCC 17978 when infected with PhAb24 also showed no sign of upregulation (Supplementary Fig. 8c). It is likely that DdaA does not respond to solo phage infection if not triggered on by DNA damage, but further verification is needed in future study.

## Discussion

Global DDR systems are organized as regulons by coordinating the expression of proteins involved in numerous cellular processes in response to DNA damages, such as cell division, error-prone replication, and excision repair, ensuring the recovery and survival of cells^9,36^. Among the three known global DDR systems in bacteria, namely SOS^9^, PprI-DdrO^10^, and PafBC^11^, the SOS response system is the most widely distributed one. In this study, we identify the fourth global DDR system in the family *Moraxellaceae* regulated by a WYL-domain containing protein DdaA, which acts as a transcriptional activator for genes in its regulation network (Fig. 3c-f). In evolution, the DdaA system replaced the SOS system at the common ancestor of *Fluviicoccus*, *Acinetobacter*, *Psychrobacter*, and related genera of *Moraxellaceae* (Fig. 2a). Whereas the three previously discovered bacterial DDR systems control DNA damage repair genes solely, DdaA is responsible for the activations of both canonical DDR genes and anti-phage defense genes in *Moraxellaceae* (Fig. 2). Knockout of DdaA led to impaired transcriptions of downstream genes and increased mortality of *Acinetobacter* strains in exposure to DNA damage agents and phage infections (Fig. 3 and 4). Specifically, DdaA is critical in enhanced protection against phage infection in response to DNA damage conditions (Fig. 4).

The dual function of the DdaA regulatory network may provide a selective advantage for *Moraxellaceae* bacteria in natural microbial communities (Fig. 5). Many microorganisms secrete DNA damage chemicals such as MMC^37^, colibactin^20^, bleomycin^38^ and many antibiotics^6^ to compete with cohabitating bacteria. These chemicals do not only directly cause DNA damage but also activate lytic replication of prophages in bacteria through many potential mechanisms including one intensively studied for phage lambda^39,40^. In normal conditions, phage repressor protein CI binds to the promoter of prophage genes to suppress their transcriptions. When DNA damage is caused by MMC, for example, the activated bacterial RecA protein digests LexA and triggers the SOS system for DDR. As prophage repressor CI is a homolog of LexA, RecA also digests the CI repressor as a side effect and consequently activates the lytic cycle of prophages^41^. The outbreak of phages in the microbial community^20^ enhances the antimicrobial effect of DNA damage agents causing massive death of bacteria – not only of those encoding these prophages but also of their close or distant relatives infected by broad-host phages^20^. As a unique DDR system, DdaA in *Moraxellaceae* regulates both DDR and anti-phage defense genes, thus providing more comprehensive protection against this dual effect of DNA damage chemicals^20^ (Fig. 5).

**Fig. 5.**
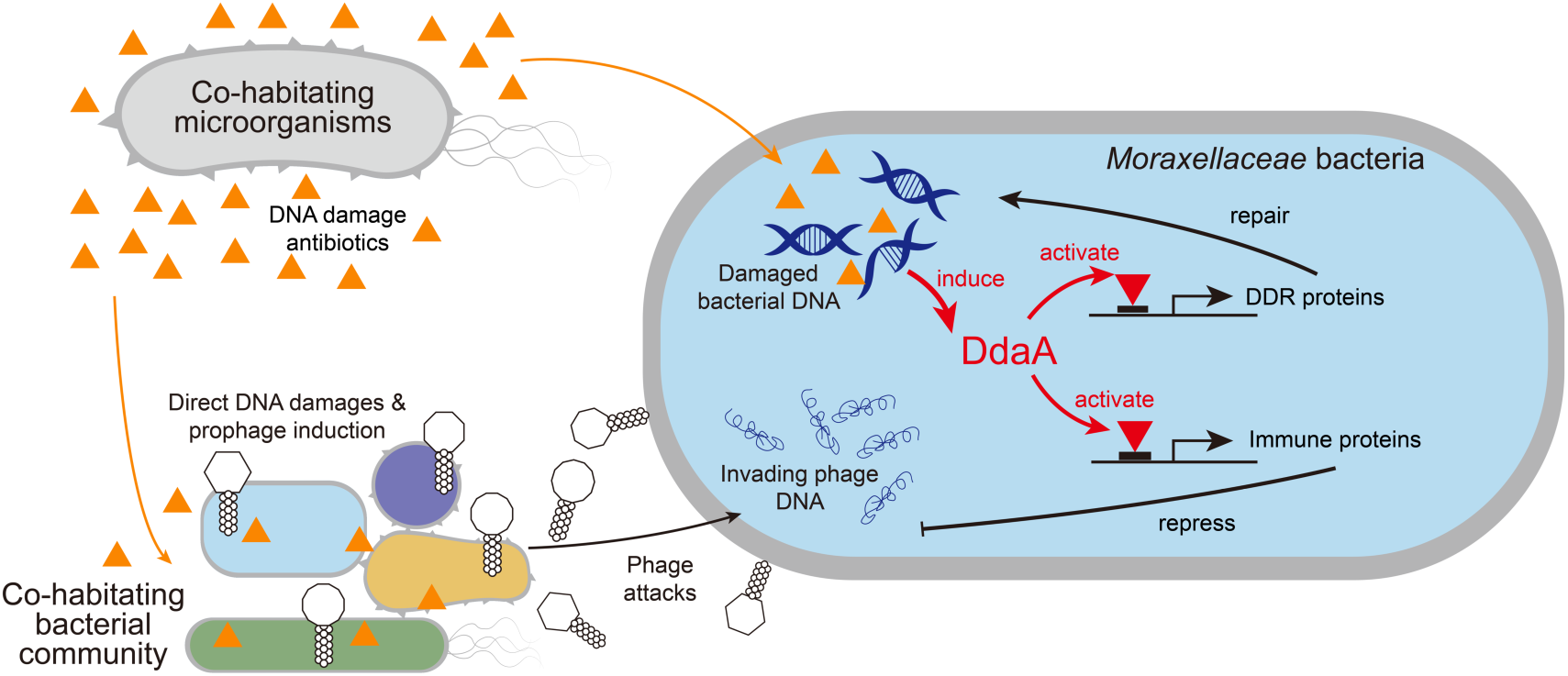
| A schematic picture showing the dual regulation mechanism of DdaA in DDR and anti-phage defense of *Moraxellaceae* under the stress of DNA damage antibiotics. Many microorganisms secrete DNA damage antibiotics to compete with cohabitating bacteria. These antibiotics not only cause DNA damage in the bacterial cells but also induce prophages entering the lytic cycle and consequently cause an epidemic of phages in the microbial community, which brings another threat. In many species of *Moraxellaceae*, DdaA potentially senses the DNA damage signals and activates the transcriptions of both DDR and anti-phage defense (immune) genes. The former ones repair direct cellular damages such as interstrand crosslinks and double-strand breaks, whereas the latter ones target invading phages, thus providing more comprehensive protection against DNA damage antibiotics compared to other known DDR systems.

This DdaA functioning model implies the insensitivity of *Acinetobacter* strains to certain antibiotics, phages, and their combinations. The interactions between phages and antibiotics in phage therapy can be complex and we are just beginning to uncover the behind mechanisms^42,43^. In recent clinical cases of phage therapy, phage treatment had been selected for increased sensitivity to antibiotics of multidrug-resistant bacteria^44^. Specifically, bacterial populations escaping phage infection were achieved by altering or losing their cell-surface structures such as the capsule or a drug efflux pump, which are also receptors for phage attachment. Such changes typically result in deficiency in blocking the entry of exogenous antibiotics or in exporting entered antibiotic molecules, demonstrating a genetic trade-off between phage evasion and antibiotic sensitivity. In contrast, the DdaA system identified here in *Moraxellaceae* provides a distinct mechanism of antibiotic-phages interaction (Fig. 5), that bacteria can simultaneously maintain their resistances to DNA damage antibiotics and phages in selection. When antibiotics such as ciprofloxacin and phages are applied in combination, strains with efficient DDR and phage-defense systems co-regulated by DdaA will be selected, implying a major clinical concern when treated with bacteria of this family^45,46^.

In summary, DdaA is a transcriptional activator protein that belongs to a monophyletic cluster of WYL domain-containing proteins and controls both DDR and phage inhibition in the family *Moraxellaceae*. This unique mechanism in DDR provides *Moraxellaceae* strains potential competitive advantages by resisting to both DNA damage chemicals and the following phage outbreak in environmental or hospital settings. Our finding reveals a previously unexpected challenge in combating the outstanding resistance of *Moraxellaceae* bacteria, but it also sheds light on the development of new therapeutic strategies by developing drugs specifically despairing DdaA, which are not encoded by human cells^31,47^.

## Materials and Methods

### The identification of the binding motif in *Acinetobacter*

Potential DDR genes predicted by the previous study^17^ (Supplementary Table 1) were downloaded from NCBI Entrez^48^ with Biopython script^49^. A pipeline ‘Novel-Sample’ was developed to search for the potential binding motifs. DIAMOND (blastp -b10.0 -k500 -p 16) was used to search the protein homologs of each gene against the protein sequences of 88 representative *Acinetobacter* species. The −200 to 0 regions from the translation start site of the genes of the founded homolog protein sequences were extracted by the Biopython script, concatenated into one file, and input to MEME^50^ (-dna -minw 12 -maxw 25 -mod zoops -evt 1e-20 -pal -p 32 -nmotifs 3).

### Bacterial strains, growth conditions and genomic sequencing

*A. radioresistens* ATCC 43998 was obtained from the Shanghai Bioresource Collection Center. The phage PhAb24, bacterial strains *A. baumannii* ATCC 17978, ATCC 19606, and ZWAb067 were obtained from laboratory stock. *A. baumannii* ZWAb067 was isolated from the brain tissue of a dead duck embryo, and its genomics sequence can be found in the NCBI BioProject: PRJNA1146946. *Acinetobacter* strains were grown at 37°C in Luria-Bertani (LB) broth. The working concentration of kanamycin was 50 μg/mL, and the working concentration of apramycin was 100 μg/mL.

*A. baumannii* ZWAb067 genomic DNA was extracted using FastPure Bacteria DNA Isolation Mini Kit (DC103-01), purchased from Vazyme Biotech Co. Genomic DNA was sequenced using Illumina NovaSeq PE150 (Illumina, San Diego, CA) at the Beijing Novogene Bioinformatics Technology Co., Ltd. Sequencing reads were assembled into a complete sequence using Unicycler^51^ release 5.0.

### DNA pulldown assay

Cultures of *A. radioresistens* ATCC 43998 and *A. baumannii* ATCC 17978 were grown in 500-mL broth until the OD_600_ reached 1.0. Cells were harvested by centrifugation at 8,000g for 5 min. Supernatants were discarded and cell pellets were resuspended in 40 mL wash buffer (20 mM Tris-HCl pH 7.5, 2 mM EDTA, 150 mM NaCl, 0.5% TritonX-100). 2 mL aliquots were collected in 2-mL EP tube by centrifugation at 8,000g for 5 min. Cell pellets were then lysed by B-PER® Bacterial Protein with instruction provided by Thermofisher.

The 3’biotin-TEG labeled ssDNA and the complementary ssDNA were synthesized by Sangon Biotech (Shanghai), based on the promoter region of the *recA* gene of *A. radioresistens* ATCC 43998. Two strands of DNA were annealed in an annealing buffer (30 mM Tris-HCl pH7.5, 100 mM KCl) through thermal annealing from 90°C to 4°C. 20 μL of 10-μM biotin-TEG labeled dsDNA was incubated with 50 μL MyOne™ Streptavidin C1 for 15 min. The DNA sequences are shown in Supplementary Table 10.

400 μL Cell lysate and DNA-coated beads were mixed. When purified DdaA is needed for co-factor pulldown, 15 μg DdaA was added to the solution. After gentle mixing for 30 min, beads were collected by magnets and washed with 500 μL wash buffer for five times. Finally, the beads were resuspended with 30 μL wash buffer and then analyzed by routine SDS-PAGE analysis. The Fast Silver Stain Kit (Beyotime Biotechnology, China) was used for silver stain.

### Mass spectrometry

The target protein band was cut from the gel and the peptides were extracted after enzymatic digestion. They were then analyzed in mass spectrometry Q-Exactive at the Hangzhou Precision Medicine Research Center (Hangzhou). Thermo Scientific Proteome Discoverer 2.1 was used for protein identification.

### Protein expression and purification

*ddaA* genes from *A. radioresistens* ATCC 43998, *A. baumannii* ATCC 17978, *P. aquimaris* strain KCTC 12254, *H. influenzae* strain 65290_NP_Hi3, *C. arsenatis* strain DSM 11915, *G. subterraneus* strain Red1 were cloned or synthesized by Beijing Tsingke Biotech Company, and then inserted into the plasmid vector pET-28a backbone (Supplementary Table 11). A 6×His-tag was added at the 5’-end of *ddaA* for affinity chromatography. The vectors were transformed into *E. coli* BL21 (DE3) cells for protein expression. The expression cells were grown in LB broth at 37°C, and when the OD_600_ reached 0.6, isopropyl-β-D-thioga-lactopyranoside was added to a final concentration of 0.2 mM. After another five hours of incubation, the cells were harvested by centrifugation at 8,000g for 5 min and washed with His-A buffer (150 mM NaCl, 20 mM Tris-HCl pH 7.5, 5% w/v glycerol). The cells were resuspended with 40 mL of His-A buffer and lysed by sonication, followed by centrifugation at 16,000g for 30 min to remove cell debris. The supernatant was applied to an AKTA Purifier system and loaded onto a HisTrap HP column after equilibration with His-A buffer. After washing with His-B buffer (150 mM NaCl, 20 mM Tris-HCl pH7.5, 5% w/v glycerol, 50 mM imidazole), the protein was finally eluted with His-C buffer (150 mM NaCl, 20 mM Tris-HCl pH7.5, 5% w/v glycerol, 250 mM imidazole). If desalting was needed for downstream experiments, the buffer was changed back to His-A by using the HiPrep 26/10 Desalting column.

### EMSA experiment

Site mutants of the *recA* promoter sequence from *A. radioresistens* ATCC 43998 labeled with 3’FAM were synthesized by Sangon Biotech (Shanghai) and annealed in an annealing buffer (30 mM Tris-HCl pH7.5, 100 mM KCl) through thermal annealing from 90°C to 4°C. 20 μL 50-nM 3’FAM-labeled DNA and 5 μL 4-μM DdaA and its homologs were mixed and incubated for 15 min. 5 μL mixture was loaded to 12% Native-PAGE and electrophoresised at 140V for 30 min. The gels were imaged in fluorescence mode (FAM) on Typhoon FLA 9500 (GE). The DNA sequences are shown in Supplementary Table 10.

### Phylogenetic relationship between DdaA and other studied WYL transcriptional regulators

We built a protein BLAST database based on all completed Refseq bacterial genomes in January 2023. We run PSI-BLAST^52^ (E-value 1e-60, iterates until convergence) against the database to find DdaA, PafB, PafC, DriD, BrxR, and CapW homologs. The protein sequences with < 260aa or > 400aa were removed, as DdaA of *A. baumannii* ATCC 17978 has a length of 335aa, PafB of *M. tuberculosis* H37Rv has a length of 332aa, PafC of *M. tuberculosis* H37Rv has a length of 316aa, DriD of *C. vibrioides* ATCC 19089 has a length of 331aa, CapW of *P. aeruginosa* PA17 has a length of 299aa and BrxR of *Acinetobacter* sp. NEB 394 has a length of 288aa. The homologous sequences were aligned using mafft^53^ v7.490 with the default parameter, and trimmed using trimAI^54^ v1.4 with the default parameter. The phylogenetic tree was generated using FastTree^55^ 2.1.11 with the default parameter. The tree was visualized in iTOL^56^ v5.

### Evolutionary analysis of the DdaA DDR system

In total, 402 genome sequences from the family *Moraxellaceae* and its closely related species were selected for the evolutionary analysis (based on GTDB^57^ release214, Supplementary Fig. 3, Supplementary Table 12). The genome sequences were annotated by Prokka^58^ 1.14.6. The protein orthologous groups were determined by Orthofinder^59^ 2.5.5 with default parameters. The MSA of the 120 marker genes provided by the GTDB was used for the phylogeny analysis of the 402 genomes. BMGE^60^ 1.12 was used to trim the MSA with default parameters. The phylogenetic tree was generated using FastTree^55^ 2.1.11 with default parameters.

To determine the DdaA orthologues, we conducted PSI-BLAST on NCBI using the DdaA from *A. baumannii* sequence against the nr_clustered database (re-analyzed on Jan.22^nd^, 2024). The ‘Expect threshold’ and ‘PSI-BLAST incl. threshold’ were set to 1e-60, The ‘Max target sequences’ parameter was set to 20,000, and the ‘word size’ was set to 3. Highly similar sequences were removed by CD-HIT (-c 0.8 -T 6 -n 2 -d 50). All Orthofinder-determined DdaA ortholog (DdaA-1 hereafter, Supplementary Text) sequences were also added to the PSI-BLAST result to perform phylogeny analysis. The protein sequences with < 260aa or > 400aa were removed, as the DdaA from *A. baumannii* has a length of 335aa. The sequences were aligned using mafft v7.490 with the default parameter and trimmed using trimAl with automated1 mode. The phylogenetic tree was generated using FastTree^55^ 2.1.11 with the default parameter. The tree was visualized by iTOL^56^ v5.

To reconstruct the transferring of DdaA genes in the evolutionary history of *Moraxellaceae*, the protein sequences of DdaA-1 were aligned by mafft v7.490, and trimmed by BMGE 1.12 with the default parameter. The phylogenetic tree was generated using IQ-Tree^61^ multicore version 2.0.3. The Best-fit model found was LG+R6. The support value of the phylogenetic tree was analyzed with the ultrafast bootstrap (UFBoot) feature 5,000 times. ALEml_undated algorithm of the ALE^62^ package was used to determine the duplications, losses, intra-LGT (transfer within the sampled genome set), and originations event of DdaA-1. Species tree of 402 genomes and DdaA-1 bootstrap trees generated above were used as input. The result was summarized by ALE helper (https://github.com/Tancata/phylo/tree/master/ALE).

### Identification of genes that are regulated by DdaA, LexA, or UmuD

The promoter sequences were defined as −200 to 0 bp from the translation start site. Biopython script was used to extract the promoter sequences. As DdaA binding motifs reside in the −350 region, the −400 to 0 region was manually extracted as *zorA* promoter. All genes encoded in 402 genomes were searched by the FIMO program^63^. The hit with a *P* value lower than 5e-6 was considered to have a DdaA binding motif in the promoter, indicating that DdaA regulates the gene’s transcription. The same process was applied to find genes that are regulated by LexA and UmuD proteins.

### CRISPR and Phage determination

The CRISPR-Cas system was detected by CRISPRCasFinder^64^ 4.3.2, and the spacer sequences were extracted from result files. The prophage sequences were detected by Phispy^65^ 4.2.6. BLASTN program was used to find the spacer and target phage pairs. To find more potential spacer-phage pairs, we did not restrict that the target phage DNA must have a PAM sequence. Spacer sequences were used as a query, and all phage DNA sequences were used as the subject. Only the result with no more than 1 mismatch was considered a true spacer-target pair (Supplementary Table 9).

### The generation of *ddaA*-knockout and compensation strains

The DdaA of *A. baumannii* ATCC 17978 was knocked out with a pT18-sacB-based knockout system following a similar method as introduced by Amin et al.^66^. In short, the upstream and downward sequences (around 500bp, respectively) of *ddaA* (locus tag: HKO16_RS19275) from *A. baumannii* ATCC 17978 were cloned into the *sacB*⁺ suicide vector pT18-sacB (with tetracycline-resistant gene) through In-Fusion^®^ technology, to make *ddaA*-knockout vector pT18-*sacB*-*ddaA*. The product of the In-fusion cloning reaction was transformed into *E. coli S17-1 λpir* chemically competent cell. The colonies grown on the tetracycline LB plate were selected, and the successful transformation was confirmed by sequencing PCR products using M13-F and M13-R primers. *A. baumannii* ATCC 17978 and *E. coli S17-1 λpir*:: pT18-sacB-ddaA were grown in LB broth with and without Tc antibiotic, respectively, till OD_600_ reached 0.8. The 700 μL *A. baumannii* culture was mixed with 700 μL *E. coli culture,* and the mixed culture was incubated at 37°C overnight. The culture was spread on a tetracycline+ampicillin LB plate and grew at 37°C overnight. The colonies on the plate were selected and transferred to liquid LB broth containing 10% sucrose for *sacB* selection. Successfully knockout of *ddaA* was validated by DNA sequencing (Supplementary Fig. 7a). To obtain the complemented strain, PCR-amplified wild-type *ddaA* with its promoter region was cloned into the plasmid pWH-1266 and transformed into the *ddaA*-knockout strain.

The DdaA of *A. baumannii* ZWAb067 was knocked out with CRISPR-Cas-based strategy^67^. The specially designed sgRNA expression vector pSGAb-km and ssDNA were transferred to the ZWAb067::pCasAb-apr electrocompetent cells (Supplementary Table 10). The knockout of *ddaA* was confirmed by Sanger sequencing (Supplementary Fig. 7a), and the pCasAb and pSGAb vectors were removed with *sacB* selection.

### Phenotyping of DNA damage treatments

Wild-type *A. baumannii*, the *ddaA*-knockout strain, and the complemented strain were resuscitated on LB plates and grew overnight at 37°C. The fresh colony was picked up into liquid LB broth and grew at 37°C till OD_600_ reached 0.6. Then, each 1 mL culture was transformed into a 1.5-mL EP tube and centrifuged at 4,000g for 5 min. The supernatant was then discarded. To test viability, the cells were resuspended with LB broth containing various concentrations of MMC or irradiated with by CL-1000 Ultraviolet Crosslinker. After incubation at 37°C for 30 min, the cell culture was placed on an LB plate and grew overnight at 37°C.

### Transcriptome sequencing and RT-qPCR

*A. baumannii* strains were resuscitated on the LB plate. The fresh colonies were picked up into liquid LB broth and grew at 37 °C till OD_600_ reached 0.6. Each 2 mL culture was transformed on a clean glass surface. The UV radiation group was treated with 15 J/m^2^ of UV radiation when testing DNA damage response or 5 J/m^2^ when testing anti-phage defense ability, while the control group was held at room temperature. As for MMC related experiments, the fresh colonies were picked up into liquid LB broth and grew at 37 °C till OD_600_ reached 0.6. The MMC was added to the experimental group culture at final concentration of 1 μg/mL for testing DNA damage response or 0.5 μg/mL for testing anti-phage defense ability. Lower doses of UV or MMC were used for testing anti-phage defense ability, to minimize the inhibition effect on the growth of bacterial cells. The cultures were then incubated at 37°C for 15 min. Then, the culture was immediately frozen in liquid nitrogen and stored at −80°C. All RNA sequencing and RT-qPCR experiments were conducted in three biological replications.

Total RNA was extracted using the RNAprep Pure Cell/Bacteria Kit RNAprep Pure, DP430, and purified using the RNAClean XP Kit (Beckman Coulter). The remaining DNA was digested using the RNase-Free DNase Set (Qiagen). Libraries were constructed using U-mRNAseq Library Prep Kit AT4221, KAITAI-BIO) with Ribo-off rRNA Depletion Kit (Bacteria) (Vazyme) and then sequenced at the Illumina NovaSeq platform to generate 150-bp paired-end reads in KAITAI-BIO company. Sequencing reads were processed using TrimmomaticPE^68^ and mapped to the representative genome of *A. baumannii* ATCC 17978 (RefSeq accession: GCF_014672775.1) using HISAT2^69^. The expression matrix was calculated using StringTie^70^ and DESeq2^71^ to determine the differentially expressed genes.

For the RT-qPCR experiment, total RNA was extracted using the TransZol Up Plus RNA Kit. Complementary DNA (cDNA) synthesis was performed using HiScript III RT SuperMix kit (Vazyme) according to the manufacturer’s protocol. Briefly, the purified RNA was added into an RNase-free centrifuge tube containing a gDNA wiper mix. The mixture was incubated at 42°C for 2 min to remove genomic DNA. Then, a 5× Hiscript III enzyme mix was added to the mixture of the previous step. After mixing, the reverse transcription reaction was performed at 37 °C for 15 min and then at 85 °C for 2 min. The cDNA is used as the template for RT-qPCR with gene-specific primer. Agilent Mx3005P automatically generates the Ct value. *rpoB* was used as the reference gene. The primer sequences for RT-qPCR are shown in Supplementary Table 10.

### Phage purification

The phage PhAb24^72^ stock was added to 5 mL of exponentially growing wild-type *A. baumannii* ATCC 17978. Cultures were incubated overnight at 37°C. Supernatant from overnight culture was then filtered through a 0.22-µm membrane and stored at 4°C. The titer of PhAb24 was determined by the soft-agar overlay assay using the wild-type ATCC 17978.

### The growth assays and plaque assays for testing *A. baumannii* strains’ resistance to phage infection

We generate a series of strains originating from *A. baumanii* ATCC 17978 harboring the RM system from *A. baumanii* ATCC 19606. To preserve the gene neighbor of the RM system, we inserted genes from ATCC 19606 defense locus6 (PADLOC-DB v2.0.0) into the pSGAb backbone. The ATCC 17978 wild-type, ATCC 17978 harboring RM system (abbreviated as strain RM), strain ATCC 17978 harboring RM system with mutated DdaA binding motif (abbreviated as strain RM-mutated motif), and ATCC 17978 *ddaA*-knockout strain harboring RM system (abbreviated as strain △*ddaA*-RM) were grown at 37°C till OD_600_ reached 0.6. If needed, MMC was added to the cultures at a final concentration of 0.5 μg/mL. The cultures were incubated at 37°C for 30 min to activate the DdaA network, then 2 mL cultures were harvested by centrifugation at 7,000 g for 1 min. The supernatant was discarded, and the cell pellet was resuspended with 2 mL of free LB broth. For growth assays, 200 μL resuspended culture was transferred to the 5-mL fresh LB broth, and PhAb24 was added to the culture at multiplicities of infection (MOI) of ∼0.02. The OD_600_ was automatically recorded every 5 min using the Scientz^TM^ microbial growth curve analyzer MGC-200. Three biological replications were conducted. For plaque assays, 200 μL resuspended culture was transferred to the warmed 3-mL soft agar LB broth, and placed on LB agar forming an overlay layer. The plate was incubated at 37 °C for 15 min allowing the surface to dry. Then the serial diluted PhAb24 was placed on the surface. MMC and ciprofloxacin concentrations were 0.5 μg/mL and 0.2 μg/mL in plaque assays, respectively.

### The assays for testing exogenous DNA removal ability of *A. baumannii* native CRISPR I-F system

To perform plasmid interference assay, the pSGAb plasmid^67^ was used as the backbone and modified with PCR-based point mutation technology to insert CRISPR-Cas target sequences, forming pTarget. pTarget contains 2 target DNA sequences which are recorded by the *A. baumannii* ZWAb067 native CRISPR array. The target DNA can be found in the prophage region of *A. baumannii* strain AB046 and the prophage region of *Acinetobacter pittii* strain FDAARGOS 1397, thus we used pSG-target to mimic the invasion of prophage induced by DNA damage. The spacer information and point mutation primers are shown in Supplementary Table 10. To perform plasmid interference assay, the pControl and pTarget plasmids were diluted to 100ng/μL and 1.5 μL diluted DNA was transformed to 50 μL electrocompetent cells of wild-type *A. baumannii* ZWAb067, *ddaA*-knockout strain or complemented strains by electrotransformation. After incubation at 37°C for 1 h, the transformed cells were placed on the kana-LB agar, and the colony was manually counted after incubation at 37°C for 16 h.

We also used qPCR to test the relative abundance of pControl to pTarget in the overnight culture. The pControl and pTarget plasmids were mixed at a 1:1 ratio of 50 ng/μL final DNA concentration. 1.5 μL diluted DNA was transformed to 50 μL electrocompetent cells of wild-type *A. baumannii* ZWAb067, *ddaA*-knockout strain, complemented strain, or wild-type *A. baumannii* ATCC 17978 by electro-transformation. The transformed cells were transferred to 5 mL fresh LB broth and incubated at 37°C for 1 h. Then the kanamycin antibiotic was added to the cell cultures at a final concentration of 50 μg/mL. The cell cultures were incubated with shaking at 37°C for 16 h, and the cells were harvested by centrifugation at 16,000g for 5 min. The supernatant was discarded, and the cell pellet was resuspended with 2 mL of sterile ddH_2_O. The cell pellet was rewashed with 2 mL of sterile ddH_2_O and was resuspended with 500 μL of ddH_2_O. Then the resuspended cultures were placed in the boiling water for 10 min and used as the direct qPCR template. The qPCR primers were specially designed to distinguish pControl and pTarget and are shown in Supplementary Table 10.

## Acknowledgments

We thank Dr. Bu Xu of the Southern University of Science and Technology for helping interpret the ALE results. This research was funded by the National Natural Science Foundation of China (32370028, 32200016, 42376113, and U22A20338) and the Zhejiang Provincial Natural Science Foundation of China (LQ23C010002 and LGN22C010002).

## Author Contributions

Y.H., L.F., H.L., and Y.Yu conceived the study. S.S. and W.W. conducted bioinformatical assays. S.S., S.Z., H.L., and Y.Z. designed the experiments, constructed the *ddaA*-knockout strain, and conducted related biochemical assays. S.Z., X.H., S.C., Z.H., and L.W. performed molecular cloning and protein purification. B.T., H.X., and D.A. conducted the biological phenotyping assays. X.Z., Q.S., Y.Yao, and W.Z. cultivated and purified bacteria strains and phages. S.S., L.F., H.L., and Y.H. wrote the manuscript. All authors took part in data analysis.

## Supplementary Information

Supplementary Information is available for this paper.

## Data availability

The raw data of RNA-seq has been deposited in NCBI under the BioProject: PRJNA895108. *A. baumannii* ZWAb067 genomic sequence can be found in the BioProject: PRJNA1146946.

## Code availability

The pipeline Novel-Sample is available online at: https://github.com/songshuang1996/Novel-Sample.

