## Supplementary Information for "A regulator controls both DNA damage response and anti-phage defense networks in *Moraxellaceae*"

### Supplementary Text

#### The difference between DdaA and other WYL domain-containing proteins

Besides DdaA, five representative WYL-family proteins including PafB and PafC from *Mycobacterium tuberculosis* H37Rv, DriD from *Caulobacter vibrioides* ATCC 19089, CapW from *Pseudomonas aeruginosa* PA17, and BrxR from *Acinetobacter* sp. NEB 394 was selected for comparison[1]. The WYL and WCX domains located at the C-terminal of these proteins are considered to have a function in detecting negatively charged ligands such as ssDNA[2-4]. Distinct from other DDR regulators, the first of the two arginine residues in the β4 strand and the β4/β5 loop, which manages the ssDNA interaction[1], was substituted by alanine in DdaA, which might lead to a different regulation mode (Supplementary Fig. 1a). Moreover, Multiple sequence alignments of the HTH domains of these WYL domain-containing proteins, which are responsible for recognition and binding to the promoter motifs[1], showed that the amino acid residues for base recognition such as the two arginine residues in α3 helix were highly conserved in the DdaA homologs (Supplementary Fig. 1b).

#### The binding motifs of DdaA and DriD are similar

The DriD binding motif in *Caulobacteraceae* was reported as CGAC-N7-GTCG recently[5], while the DdaA binding motif we discovered is CGWCA-N5-TGWCG. To compare these motifs in detail, we generated the DriD binding motif (Logo plot, Supplementary Fig. 1c) using a similar method that we used to predict the DdaA binding motif. DriD-regulated proteins YP_002516524.1, YP_002516514.1, YP_002516823.1, YP_002517748.1, YP_002518517.1, and YP_002516036.1[3] were used to search the protein homologs of each gene against all protein sequences of *Caulobacteraceae*. The promoter sequences were analyzed using the MEME program.

As expected, the Logo plots of DdaA and DriD showed high similarity (Supplementary Fig. 1c). While the MSA showed that the HTH domains (which interact with the base in dsDNA) of DdaA and DriD were not conserved (Supplementary Fig.

1b), the HTH domains of DdaA and DriD had protein tertiary structural similarity (Supplementary Fig. 1d), which may explain why DdaA and DriD bind to similar motifs. This tertiary structural similarity also indicates that they may share a common ancestor.

**DdaA forms homodimer**

To investigate the oligomeric form of DdaA in solution, we conducted size exclusion chromatography. Purified *Acinetobacter baumannii* DdaA in His-A buffer was applied to Superdex 200 Increase 10/300 GL. The flow velocity of the system was set to 0.5 ml/min. The size exclusion chromatography of *A. baumannii* DdaA showed an elution volume at 13.4 mL (peak), giving a molecular mass of 93 kDa (Supplementary Fig. 2a), suggesting that DdaA in solution is at least a dimer.

The model of DdaA homologs was predicted by the local version AlphaFold-Multimer v2.3.2 on CPU using the reduced databases[6]. The databases were updated to August 2023. The predicted model showed that two monomers of DdaA of *Acinetobacter baumannii* formed a well-symmetric homodimer (pTM: 0.80792452, ipTM: 0.80098855. Supplementary Fig. 2b). Moreover, in consistency with class A/C WYL domain-containing proteins, the predicted model exhibited an N-terminal wHTH domain (light blue), a WYL domain (green), and a WCX domain (orange) (Supplementary Fig 2b).

**The evolutionary origins of DdaA homologs in *Moraxellaceae***

DdaA homologs in *Moraxellaceae* and *Alcanivoracaceae* could be divided into two groups. Most DdaA homologs in *Moraxellaceae* and *Alcanivoracaceae* formed one cluster, which we named DdaA-1. Some DdaA homologs lay far away from this cluster, which we named DdaA-other (Supplementary Fig. 4). The DdaA-1 copies were vertically inherited in *Moraxellaceae* species. DdaA-other copies in *Moraxellaceae* were horizontally transferred. These copies may recognize the same DNA motif as DdaA-1.

**The *lexA* and *umuD* homologs have a complex evolution routine in *Moraxellaceae***

The Orthofinder software has clustered the LexA and UmuD into one homologous group. Various studies have shown that only *umuD* homologs were present in the *Moraxellaceae* genomes[7-9]. Thus, we managed to separate these two groups. Through phylogenetic analysis, we found that all LexA proteins formed one clade, while other proteins formed various clades (Supplementary Fig. 6a). We named all these proteins UmuD in this study, and these proteins show complex evolutionary history (Supplementary Fig. 6b).

**The reverse-transcription quantitative PCR (RT-qPCR) results**

We selected *ddaA*, the radical SAM protein-encoding gene, dGTPase, *recA*, *uvrA*, *uvrC*, and *aciT* as candidate genes for RT-qPCR. The RT-qPCR results support the RNA sequencing outputs that the transcription of the radical SAM protein-encoding gene, dGTPase, *recA*, *uvrA*, *uvrC*, and *aciT* genes were up-regulated in the wild-type strain but not in the *ddaA*-knockout strain after UV radiation (Supplementary Fig. 7d). The transcription of DdaA remained unchanged in the wild-type strain before and after UV radiation, and this result indicates that DdaA does not influence other genes' expression by the change of its expression.

There was a significant increase in the *ddaA* transcription level in the complemented strain compared with the wild-type, which can be explained by the multiple copies of vectors per cell[10]. The higher level of the DdaA protein led to a higher basal transcriptional level of DdaA-regulated genes including the radical SAM protein-encoding gene, dGTPase, *recA*, *uvrA*, *uvrC*, and *aciT* in the complemented strain compared with the wild-type (Supplementary Fig. 7d). The transcriptional level of these genes remained unchanged in the complemented strain after UV radiation, which can be explained by the already over-expression of these genes caused by a higher expression level of DdaA (Supplementary Fig. 7d).

When testing the transcription of DdaA-regulated genes of *A. baumannii* when infected by phage. The ATCC 17978 wild-type was grown at 37°C till $OD_{600}$ reached 0.6, and PhAb24 was added to the culture at multiplicities of infection (MOI) of ~0.2. The bacterial cells were collected every 20 minutes.

### Supplementary Figures

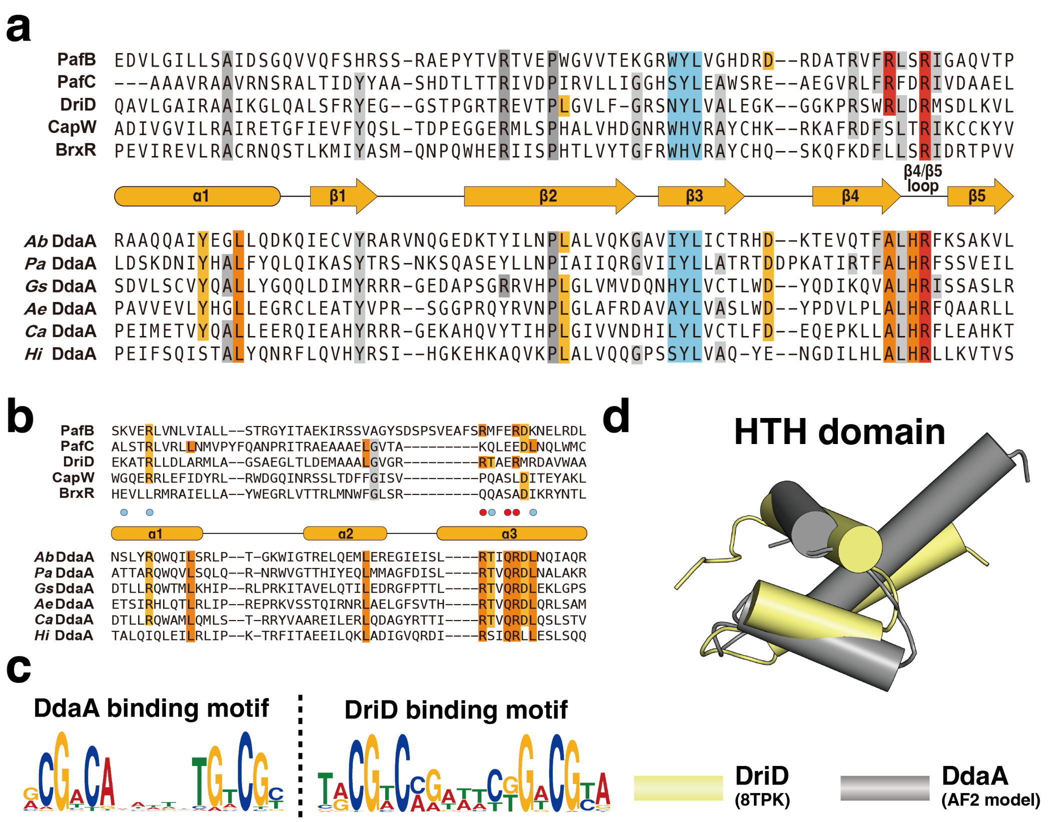


**Supplementary Fig. 1 | The comparison between DdaA and other WYL domain-containing protein. a**, Multiple sequence alignments of the WYL domains of WYL family proteins. The amino acids conserved in DdaA are colored in yellow or orange. Other highly conserved amino acids are colored in grey. The WYL motif is highlighted in blue, and the conserved residues involved in nucleotide (ssDNA) binding are highlighted in red. **b**, Multiple sequence alignment of the HTH domains of WYL family proteins. The blue dots highlight the amino acid for phosphate contacts, and the red dots highlight the amino acid for making base-specific hydrogen bonds from DriD structure[5] (PDB:8TPK). The amino acids conserved in DdaA are colored in yellow and orange based on their conservatism. Other highly conserved amino acids are colored in grey. **c**, The DdaA and DriD binding motifs. **d**, The comparison of the HTH domain structure between *A. baumannii* DdaA (predicted by AlphaFold2) and DriD (PDB:8TPK).

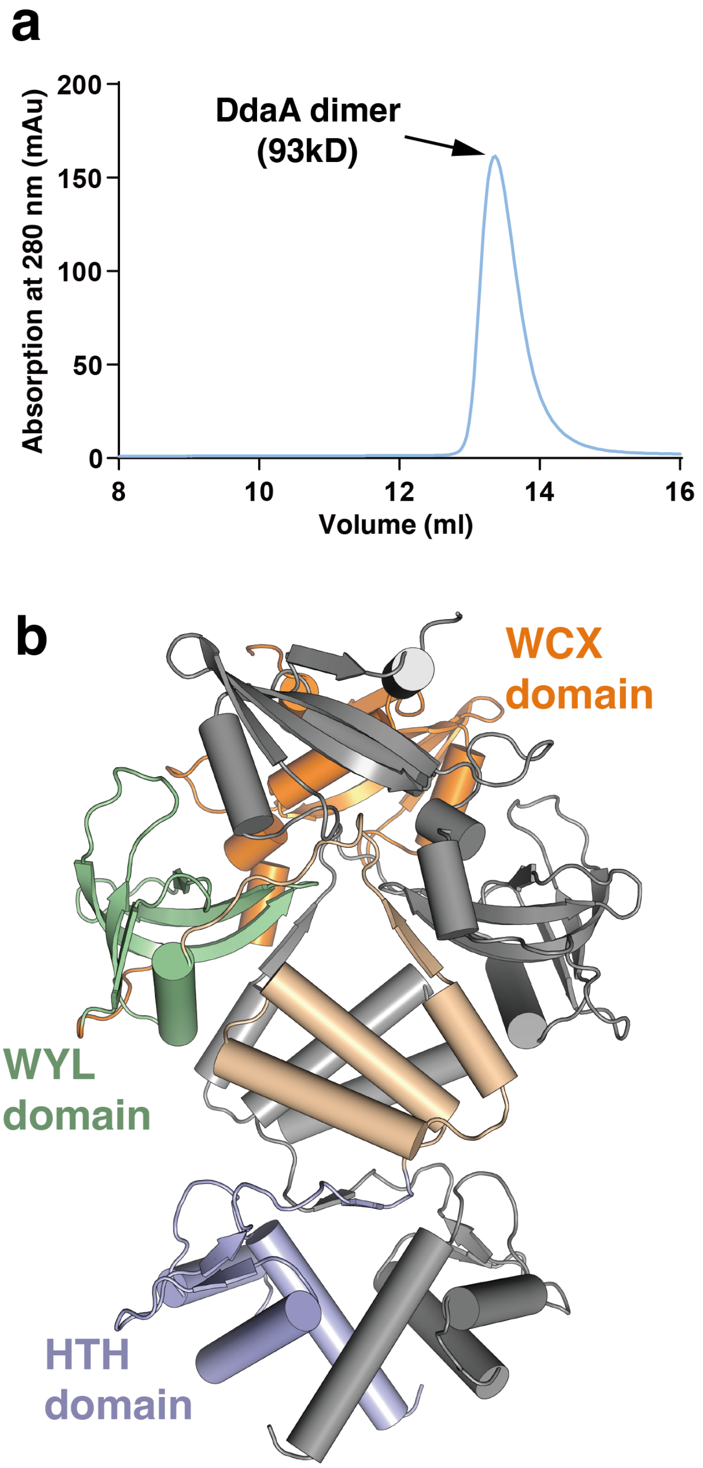


**Supplementary Fig. 2 | DdaA forms homodimer. a**, Size exclusion chromatography of DdaA from *A. baumannii*. **b**, The AlphaFold prediction of the DdaA of *A. baumannii*.

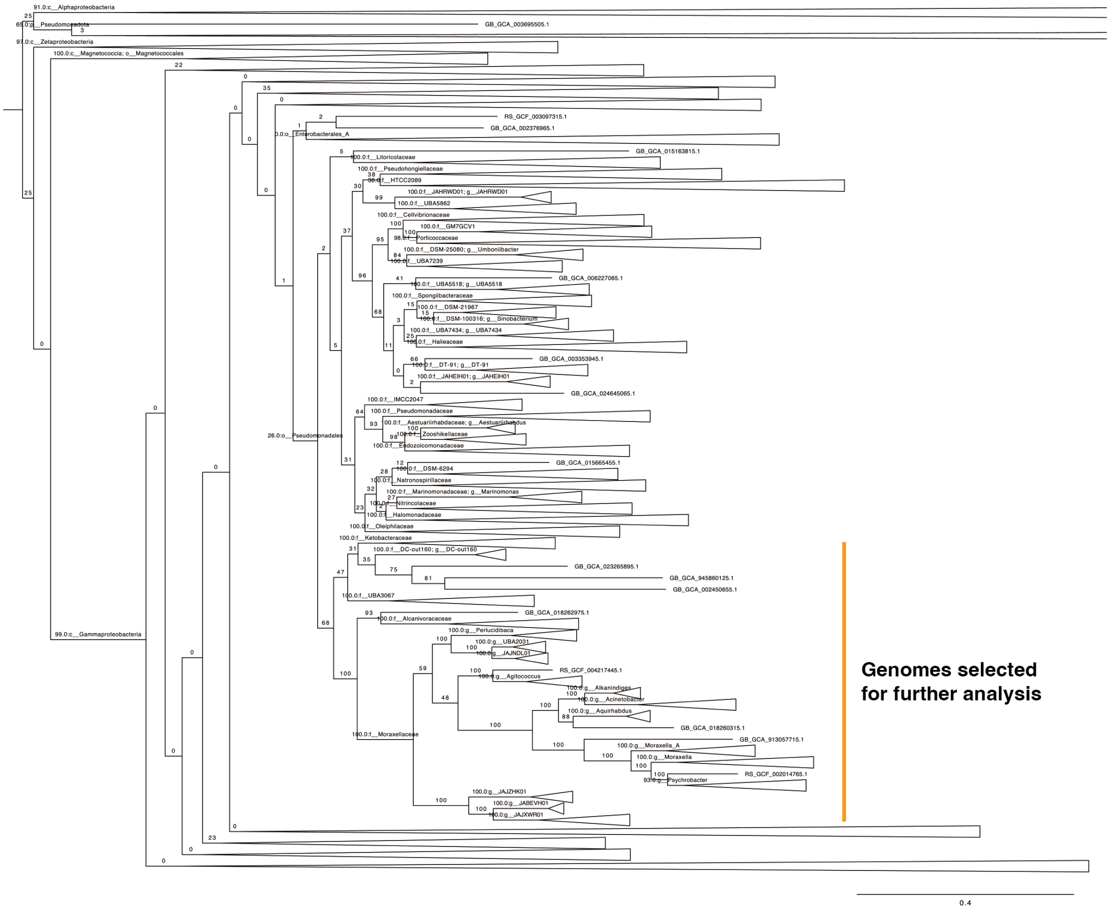


**Supplementary Fig. 3 | Evolutionary position of *Moraxellaceae* in the order *Pseudomonadales*.** The subtree is extracted from the bacterial tree of GTDB release 214. The orange bar shows the genomes used for analysis in Fig. 2 and Supplementary Fig. 6.

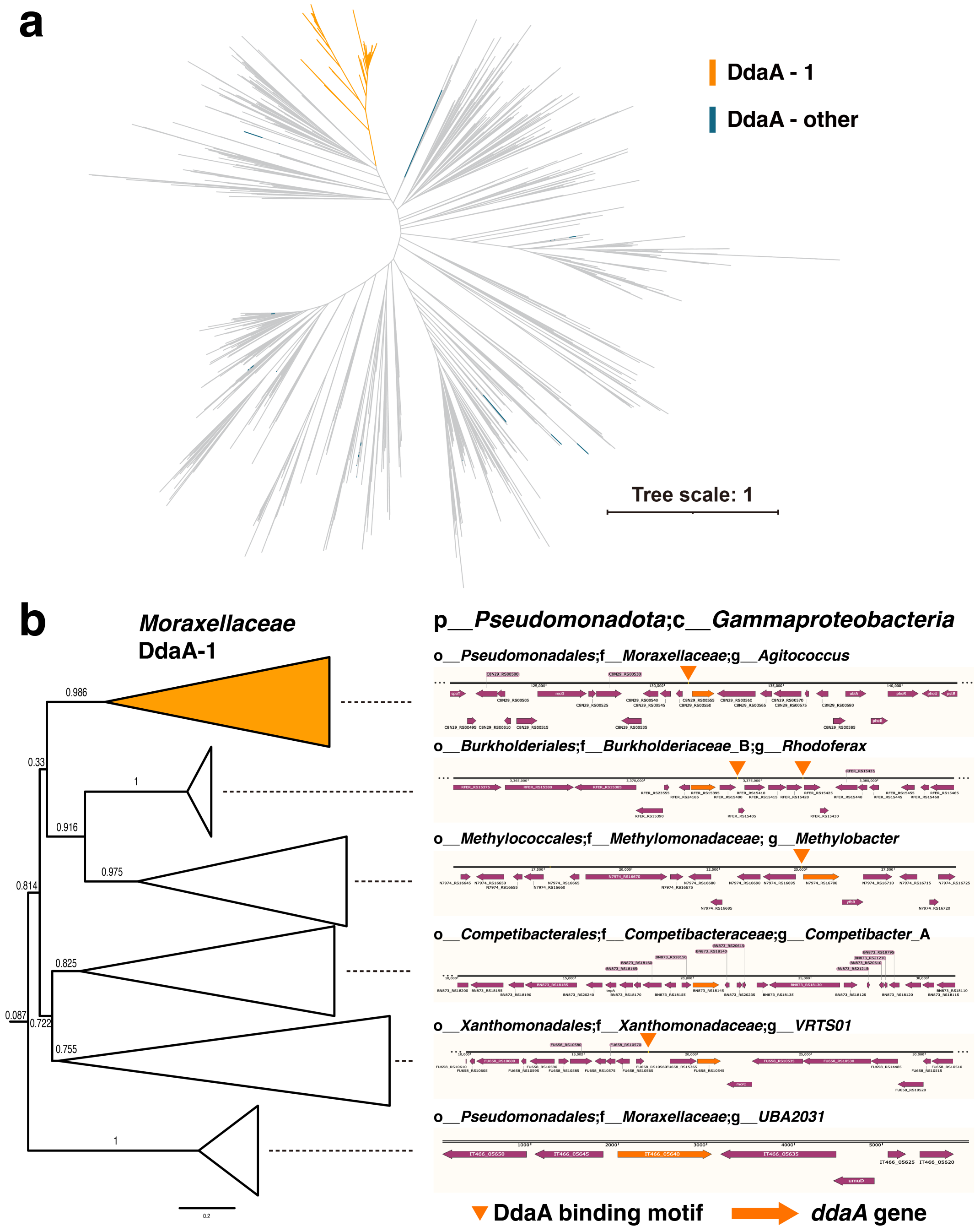


**Supplementary Fig. 4 | The phylogenomic maximum-likelihood tree of DdaA homologs in *Moraxellaceae*.** **a**, DdaA homologs in *Moraxellaceae* and *Alcanivoracaceae*. **b**, The DdaA gene tree reveals that DdaA-1 might be transformed into the common ancestor of *Alcanivoracaceae* and *Moraxellaceae* from a *gammaproteobacteria*. One *ddaA* gene and its neighborhood were selected from each cluster to show the genomic context of these *ddaA* genes.

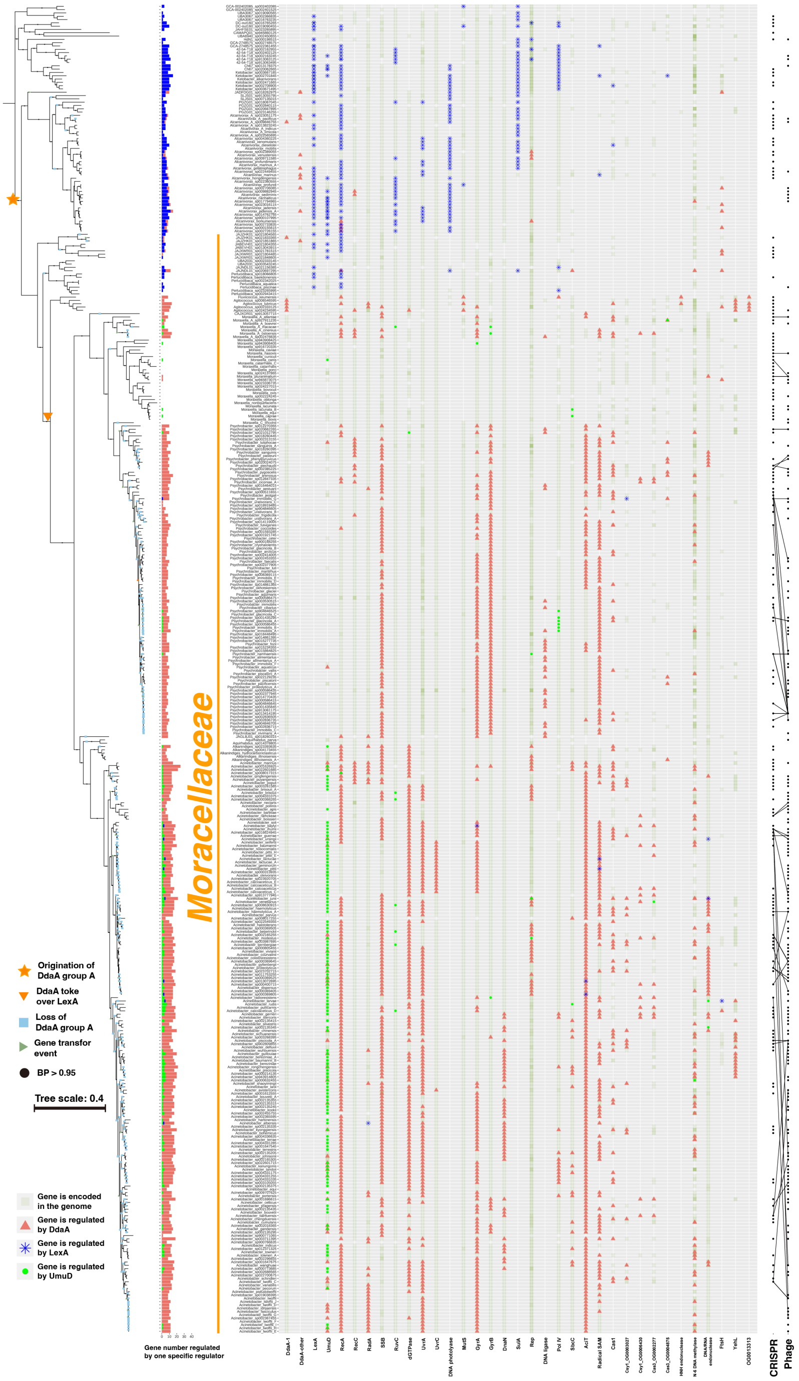


**Supplementary Fig. 5 | Detailed distribution and evolutionary route of DdaA and its binding motif in *Moraxellaceae*.** The phylogenomic maximum-likelihood tree of *Moraxellaceae* species is shown on the left, with loss and gain event prediction according to ALE analysis. The bar plot shows the number of genes regulated by DdaA in the specific species. The light green square indicates that this gene is present in this genome, the blue star indicates that LexA regulates this gene, the green circle indicates that UmuD regulates this gene, and the red triangle indicates that DdaA regulates this gene. The cross-immunity relationship of the CRISPR-Cas system and prophage is shown on the right.

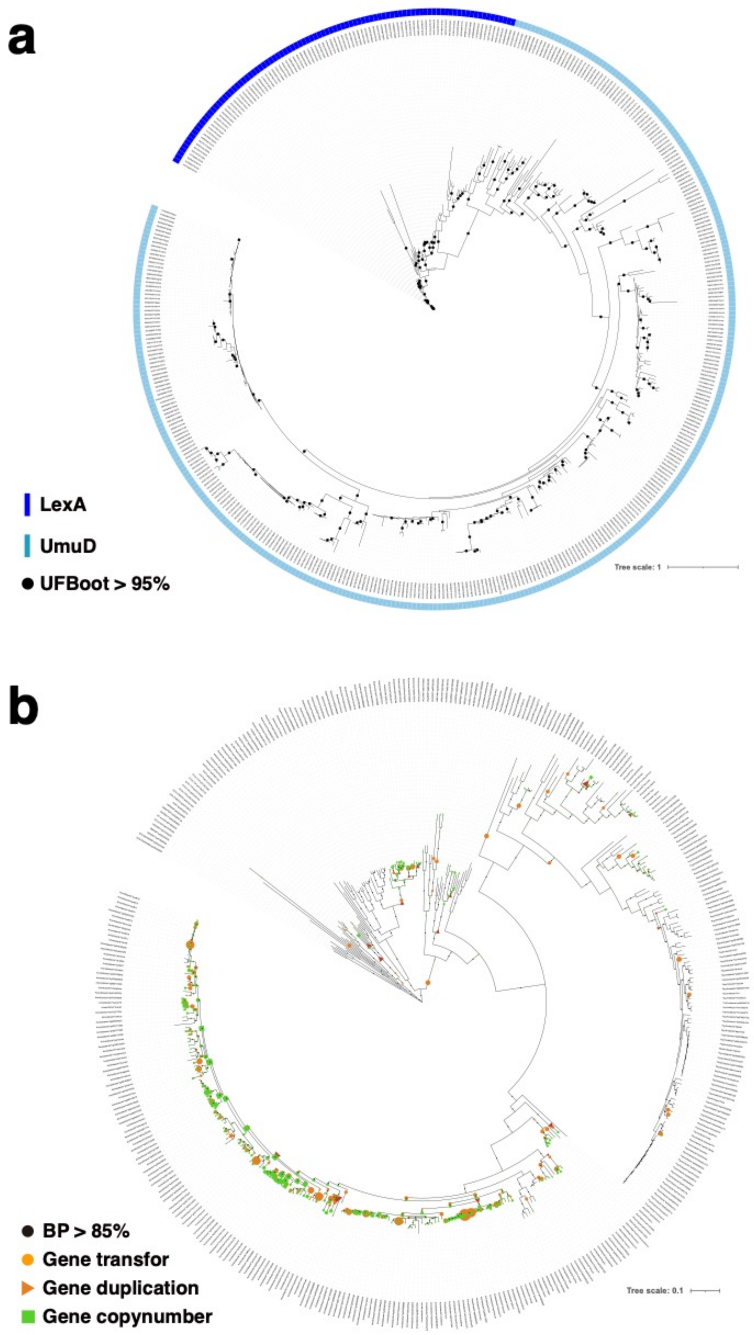


**Supplementary Fig. 6 | The phylogenomic maximum-likelihood trees of LexA and UmuD homologs. a**, UmuD and LexA were clustered into one homologous group. LexA forms one monophyletic clade, while other protein forms various clades. Our research defines all proteins outside of the LexA clade as UmuD. The label consists of two parts, Genbank ID is shown on the left side of the caret, and the locus tag provided by Prokka is shown on the right. **b**, ALE analysis result shows that the *umuD* homologs were horizontally transferred to the species in the family of *Moraxellaceae* from many events.

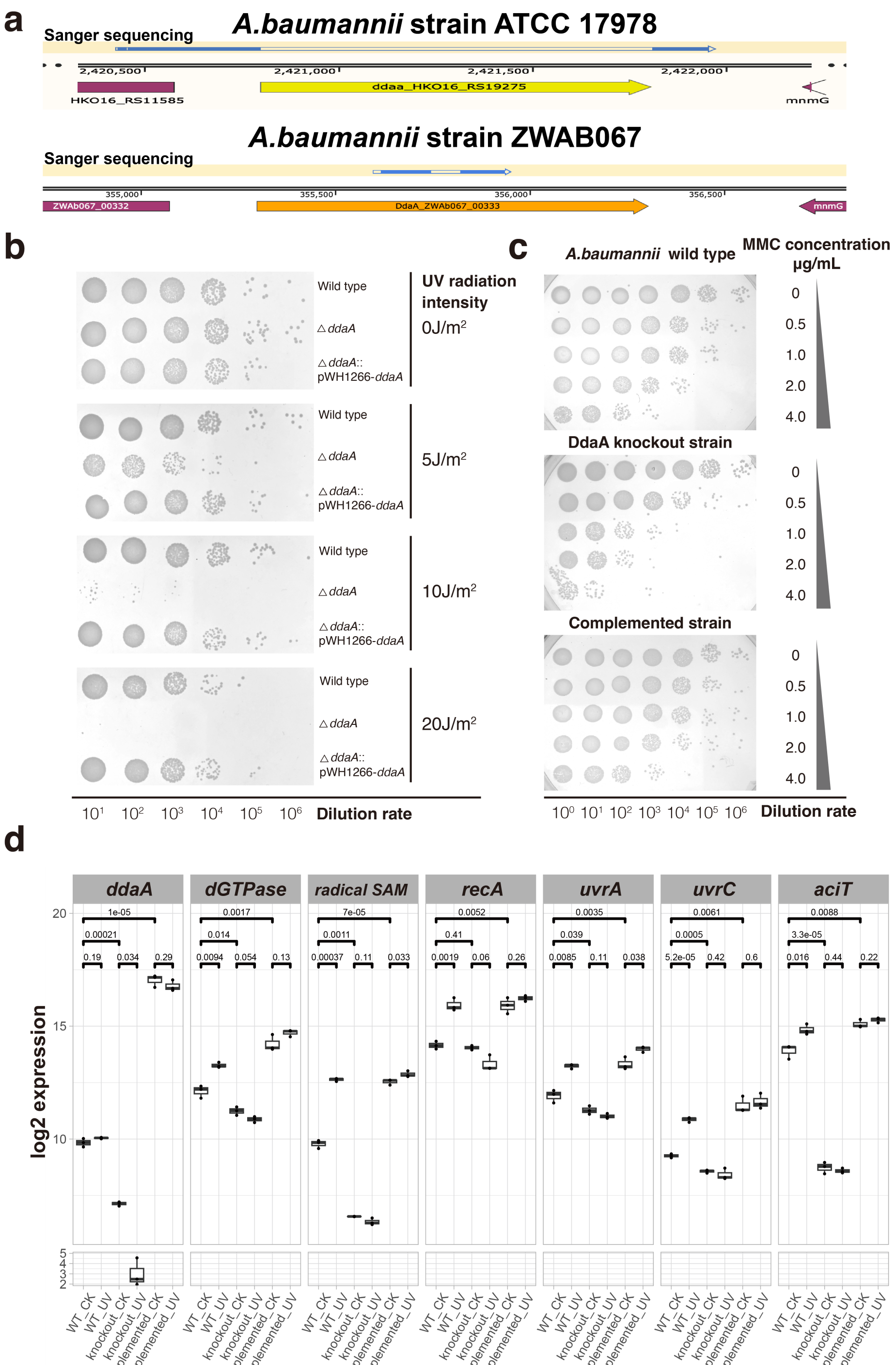


**Supplementary Fig. 7 | Knockout DdaA severely impairs bacterial stress response under various conditions.** **a**, DNA sequence, generated from Sanger sequencing, aligned to *Acinetobacter baumannii* ATCC 17978 and ZWAb067 genomes, conforming knockout of the *ddaA* gene. The blue region indicates that the sequence can be aligned, while the blank region shows that the DNA sequence has been knocked out. **b**, MMC treatment viability. **c**, UV radiation viability. **d**, The RT-PCR results show expression patterns of DdaA-regulated genes. Three biological replications were conducted.

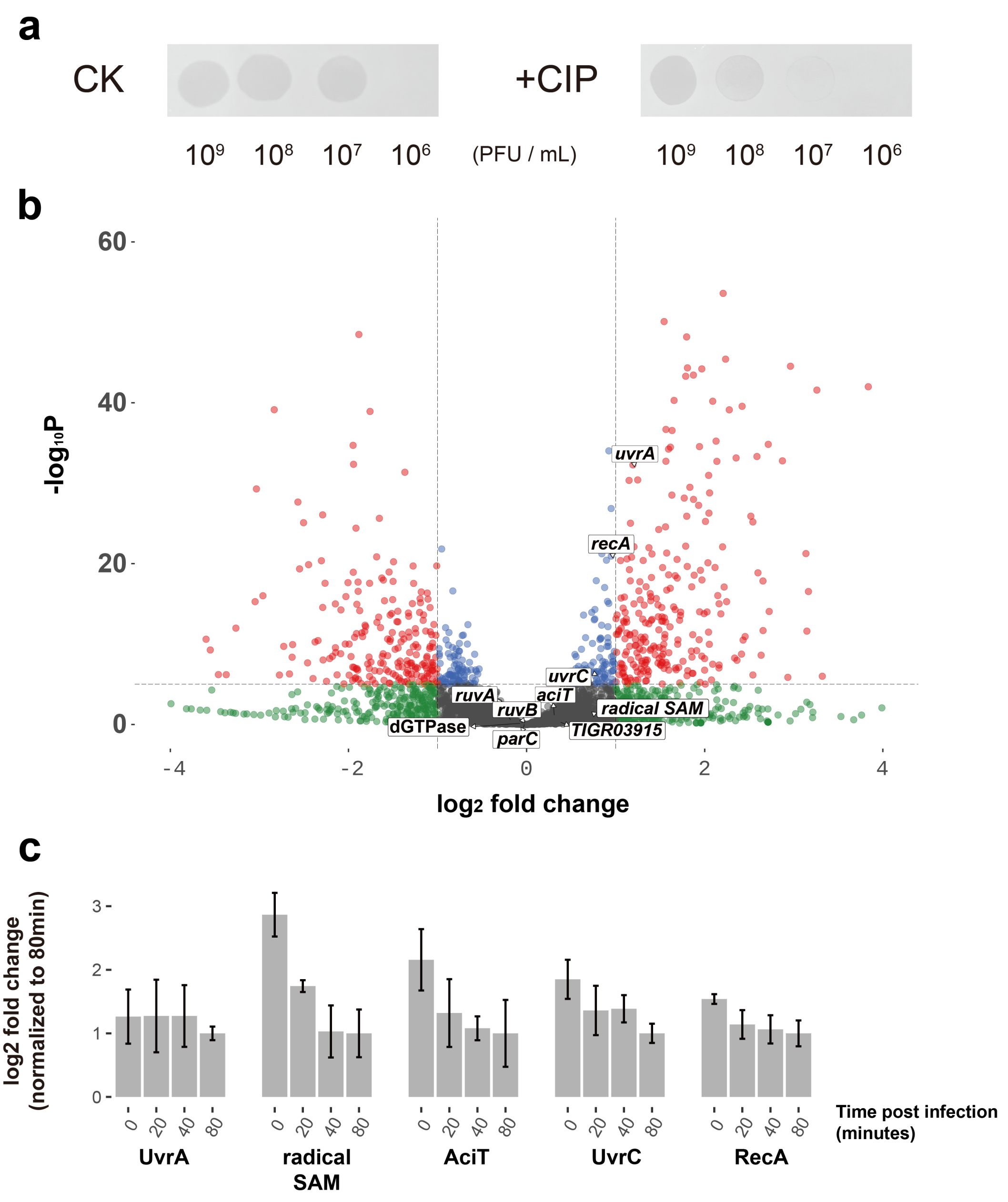


**Supplementary Fig. 8 | Ciprofloxacin treatment can enhance anti-phage defense ability, and DdaA network does not directly respond to phage infection. a,** phage plaque forming assays obtained from co-cultures of *A. baumannii* pRM and phage PhAb24 when treated with or without 0.2 μg/mL ciprofloxacin. **b**, Volcano plots showing the transcriptional level changes of genes of *A. baumannii* AB1 when infected by phage φAbp1[11] at an MOI of 10. **c**, The RT-qPCR assays result of *A. baumannii* ATCC 17978 when infected by Phab24[12] at an MOI of 0.2 at different times.
